# Comprehensive evaluation of statistical approaches for differential metaproteomics

**DOI:** 10.64898/2025.12.10.693402

**Authors:** Tjorven Hinzke, Benoit J. Kunath, J. Alfredo Blakeley-Ruiz, Abigail Korenek, Simina Vintila, Paul Wilmes, Manuel Kleiner

**Affiliations:** University of Greifswald, partner of the Greifswald Mire Centre, Greifswald, Germany; Helmholtz Institute for One Health, Greifswald, Germany; Department of Plant and Microbial Biology, North Carolina State University, Raleigh, NC, USA; Luxembourg Centre for Systems Biomedicine, University of Luxembourg, Esch-sur-Alzette, Luxembourg; Multiomics Data Science, Department of Cancer Research, Luxembourg Institute of Health, Strassen, Luxembourg; Department of Life Sciences and Medicine, Faculty of Science, Technology and Medicine, University of Luxembourg, Esch-sur-Alzette, Luxembourg

**Keywords:** Quantitative proteomics, regression, Bayesian statistics, generalized linear mixed models, edgeR, limma, microbiome, holobiont, microbial community

## Abstract

**Background:** Metaproteomics characterizes and compares molecular phenotypes of organisms in communities by comprehensively analyzing their protein expression profiles using statistical methods. However, not all statistical methods are suitable for determining differentially abundant protein groups in metaproteomic analyses. Statistical challenges in metaproteomics include: data sparsity, non-normality, compositionality, and large between-sample variability. These challenges can potentially be addressed with several data processing steps, including imputation, normalization, transformation, and selection of the appropriate statistical tests. The potential combinations of different processing methods create a complex matrix of analysis options and it is currently unclear how these combinations impact the results of statistical tests on metaproteomic data.

**Results:** To determine what data processing methods and statistical tests are best for identifying differentially abundant proteins in metaproteomics datasets, we generated a set of thirteen metaproteomic samples with known compositions, known differences, and differing levels of complexity. These defined metaproteomes address the general challenges outlined above, using various scenarios in metaproteomic data analyses. We compared over 110 different statistical analysis combination options, including regression-based tools, general statistics inference, and machine learning techniques. We found that several combinations within the frameworks of limma, edgeR, MaAslin2, custom linear and Bayesian linear models, and random forests all offer suitable evaluation options.

**Conclusions:** We highlight key recommendations for differential expression analysis in metaproteomics. Our work enables improved assessment of statistical methods for metaproteomics by establishing a framework for testing statistical approaches, including comprehensive raw mass spectrometry data and reproducible benchmarking code.

## Background

Metaproteomics is the term used for approaches that comprehensively characterize gene expression in microbiomes at the protein level [1–3]. Metaproteomics usually involves high-resolution liquid chromatography and mass spectrometry methods to identify and quantify tens of thousands of peptides that are then used to identify and quantify thousands of proteins in each sample. Based on protein abundances that differ between samples from different environments, treatments, or conditions, we can determine how metabolic and physiological processes in the microbiome respond to situations such as changing diet in the gut [4–6], ecological succession in acid mine drainage [7], organic matter availability at different depths in the ocean [8], or host-symbiont and host-pathogen interactions [9, 10].

Despite the successful application of metaproteomics to samples from many environments there is currently no consensus on how to determine which proteins are actually differentially abundant in different groups of samples. In fact, this consensus is even lacking for regular single-organism proteomics [11–14]. Additionally, it remains to be shown whether statistical tools used for proteomics, or other - omics, work as expected for complex metaproteomic analyses.

Statistical data analysis is rendered challenging by several features of metaproteomics data (please refer to the Box for details). These features and consequential challenges include: different possible protein abundance measures, i.e., spectral counts (SpC), or area for chromatographic peaks of peptides from MS ion currents (AUC), data sparsity and concomitant imputation, batch effects and non-Gaussian data distribution, and compositionality. After taking these challenges and the resulting data pre-processing steps into account, there is still the open question of the statistical test or tool for inferring differentially abundant proteins. Comparative evaluations of statistical tools for (meta-)proteomics are mostly based on the use of spike-in proteins [15–17], which, while undoubtedly useful, cannot fully capture the complexity of environmental metaproteomes. Alternatively, evaluations use experimental data without a known ground truth, and/or simulated data [11, 16, 18], meaning that their direct transferability is still unknown. Likewise, approaches for evaluating statistical tools for other omics- and microbiome techniques include mainly experimental, down-sampled, or simulated data (e.g., [19–22]). Following from this, the use of complex samples of controlled composition (mock communities or ground truth samples) for statistical method evaluation is largely missing.

Here, we generated complex defined metaproteomes with known compositions and measured them with a standard metaproteomics workflow to use as ground-truth data. We designed our defined metaproteomes to address various challenges of metaproteomics data analysis: (a) small, but biologically important, changes in protein abundances, (b) very large changes in total protein abundances of specific species in the microbiome, and (c) potential misassignment of identified proteins to the incorrect species when sequences are very similar. We measured these communities with a data-dependent acquisition (DDA) metaproteomics workflow to use as ground-truth data. We subsequently used the metaproteomic data from these defined metaproteomes to compare various data preparation and statistical inference methods using both SpC and AUC quantification methods. We tested various commonly used statistical tools used in (meta)omics and microbial community differential abundance analysis (e.g., edgeR, DESeq2, limma; [23–25]) as well as general statistical inference methods (t-test, Wilcoxon), machine learning (random forests; [26]), and Bayesian statistics-based approaches with various combinations of normalization and/or transformation approaches. This framework for testing statistical approaches can be used in the future to expand to other data acquisition approaches such as data-independent mass spectrometry (DIA).

### Box

#### Statistical terminology

##### Discrete and continuous data

Discrete data are represented as whole numbers (integers), e.g., count data. Continuous data, on the other hand are fractional and require decimals (float).

##### Compositional data

Compositional data are proportions or percentage data, or can be represented as such, i.e., data relative to a total. Proteomics mass spectrometry data are inherently compositional: for example, a cell is made up of 10 % protein A, 5 % protein B, etc. In addition to this “biological compositionality”, the mass spectrometry data acquisition introduces compositionality, because there is an upper limit to the amount of data that can be acquired. It has been advocated that specific methods have to be used to address this feature of compositionality [27].

##### Data sparsity and imputation

Especially in complex metaproteomic datasets, not all proteins present in the sample, and certainly not all peptides of specific proteins, will be detected in all conditions, especially when proteins are lowly abundant. Some statistical procedures cannot deal with missing data or zeros, e.g., because they are using a log transformation (and the log of zero is not defined). Imputation, i.e., replacement of missing values or of zero by some specific value (or values, in the case of multiple imputation), can be used to address missing values. Multiple different imputation methods exist, ranging from replacement with a small constant (e.g., the smallest number possible in the dataset or a fraction thereof), to complex statistical procedures, and their use depends on the assumption of whether the data is missing at random (MAR) or missing not at random (MNAR), or consists of a MAR/MNAR mixture [28]. Alternatively, imputation-free models, which take the pattern of missing values into account [29] may be used, but missingness patterns are informative only for a higher number of replicates.

##### Normalization vs Transformation

Normalization is used to make data comparable between MS runs, batches, etc. For example, if different amounts of peptides were injected for the respective MS runs, one would expect this to cause differences in protein group abundances. Consequently, these effects have to be corrected. For that, a multitude of normalization methods exist (e.g., [30]), and the method used will impact the outcome of differential expression analysis. Transformations, on the other hand, convert the data to adhere to a certain data distribution. Statistical tests often operate under the assumptions of a specific data distribution (e.g., Gaussian, Poisson, etc.). To meet these assumptions, data can be transformed (e.g., log, square root, etc.). Different transformations can be suitable for the same data, and they will impact downstream statistical inference.

##### Type I error

Detection of false positives, i.e., a variable (protein group) is determined by a test as being significiantly different between groups, whereas, in fact, it is not.

##### Type II error

Detection of false negatives, i.e., a variable (protein group) is determined by a test as not being significantly different between groups, whereas, in fact, it is.

##### Power of a statistical test

Probability for a test to detect a true significant difference.

##### Overdispersion

Overdispersion means that the variability in the data is greater than that expected by a given model. This happens for example for Poisson models: in the Poisson distribution, the mean equals the variance. But in real-life count data, the variance is often larger (overdispersed).

#### Statistical tests

##### Parametric vs non-parametric tests

Parametric tests, such as the t-test, assume that the data stem from a specific underlying distribution, e.g., that they were sampled from a Gaussian normal distribution. The “parameters” in a normal distribution would, for example, be the mean and variance, which define the exact shape of the distribution. Non-parametric tests, such as Wilcoxon’s rank-sum test, do not assume a certain distribution, i.e., they have no parameters (hence non-parametric). This generally results in lower power of non-parametric tests and thus a lower ability to actually detect existing statistical differences. The assumption of independence of the samples being compared holds true for both parametric and non-parametric tests. While samples need to be independent, correlated data (e.g., tests on the same subjects, or time series) necessitates adapted approaches explicitly taking into account these correlations (e.g., paired tests, or mixed models).

##### Generalized linear mixed model (GLMM)

At the core, GLMMs are linear regression models. They assume a linear relationship between the predictor variables (e.g., the protein group abundance) and the response variable (e.g., the environmental or experimental condition). However, the linearity assumption cannot hold true when, for example, the response is constrained between 0 and 1. Here, the “generalized” comes into play: it refers to a link function, which is a function of the response variable that is linearly correlated with the predictors – for example, the log. The “mixed” refers to the inclusion of random effects in addition to fixed effects as predictors. Random effects can be batches of samples that are sampled at a specific time point or extracted together, or subjects in the case of repeated measurements, but also proteins. Mixed models can thus inherently incorporate non-independence between observations.

##### Frequentist vs Bayesian statistics

Frequentist statistics refers to “classical” null hypothesis testing, most often done by many repetitions of an experiment, i.e., many independent replicates. Bayesian statistics, on the other hand, offer a mathematical framework to update prior knowledge with data, generating posterior knowledge. For example, if the expected range of standard errors of protein abundance measurements is known from previous experiments, this can be explicitly used as a “prior”. One difference between both (combinable) approaches lies in the interpretation of the uncertainty estimations: a frequentist 95% confidence interval holds the true population value 95% of the time when repeating an experiment under the same conditions. A Bayesian credibility interval, on the other hand, is the range in which the true value is expected to be at 95% probability.

##### Random forests

Random forests [26] are a supervised machine learning technique, which can be used to infer to which class a sample belongs (or, in the case of a quantitative response, be used for regression), and to determine which variables drive the distinction between classes. It uses bootstrapped sample subsets (i.e., samples are drawn with replacement), and for classification splits these subsets to generate nodes with as little mixture between pre-assigned classes as possible. For these splits, a random subset of variables (here: protein groups) are used. The full split is referred to as a decision tree. The final result is the majority vote of the trees. As random forests can be used for high-dimensional data, do not easily overfit and render variable importance measures, they are well suited for meta-omics datasets [31].

#### General considerations for statistics in metaproteomics

##### Protein quantification in (meta-)proteomics

Spectral counts, also known as peptide spectrum matches (Spc or PSMs), are the number of peptide fragment spectra (MS/MS or MS2) that are matched to a peptide sequence. The alternative to spectral counting is the extraction of peptide ion intensity data from the LC-MS/MS data (extracted ion current, XIC). This data can be used for intensity-based quantification by determining the maximum intensity for specific peptide ions in the XIC, or by integration of the chromatographic peak area for a specific peptide ion (area under the curve, AUC). Depending on the quantification method used either count data (SpC) or continuous data (AUC) are generated, which can necessitate different data transformations.

##### Data exploration

For (meta-)proteomics, common data exploration steps include hierarchical clustering and (N)MDS plots, to examine whether the data support the expectations, and, more importantly, the experimental design. We expect these steps to be done prior to applying any statistics, see for example [2] for details.

[End of box]

## Methods

### Design and generation of defined metaproteomes

Our defined metaproteomes were mixed at the peptide level and consisted of defined combinations of pure culture peptides with background peptide mixtures (“matrices” in Table 1). All defined metaproteomes were generated in biological quadruplicates. Also, each pure culture produced in biological quadruplicates for each condition (Table 1, Supplementary Table 1). The first of three background matrices consisted of peptides from stool sampled from mice with a conventional microbiota [5], simulating a sample with a high level of complexity. The second matrix was generated from corn roots that were grown under sterile conditions, and were therefore of low complexity. The last matrix consisted of stool sampled from gnotobiotic mice with a defined gut microbiome of 13 species. This last matrix provided a medium level of complexity. For the conventional and gnotobiotic mice samples, NC State’s Institutional Animal Care and Use Committee approved all experimental activity (Protocol # 18-034-B and 18-165-B). Neither euthanasia nor anesthesia were used in the collection of these samples.

**Table 1:** Pure cultures and matrices used for generating defined metaproteomes.

| Pure cultures |  |  |
| --- | --- | --- |
| Name | Description | Culturing conditions |
| *Thermus thermophilus* | Gram-negative bacterium; not present in mouse microbiome | High and low temperatures |
| *Chlamydomonas reinhardtii* | Eukaryote; not present in mouse microbiome | High and low light |
| *Rhizobium leguminosarum* bv. *viciae* 3841 | Gram-negative bacterium; not present in mouse microbiome, plant symbiont | Strain comparison with *R. leguminosarum* VF39 |
| *Rhizobium leguminosarum* bv. *viciae* VF39 | Gram-negative bacterium; not present in mouse microbiome, plant symbiont | Strain comparison with *R. leguminosarum* 3841 |
| *Escherichia coli* | Gram-negative bacterium; potentially present in mouse microbiome | Complex (LB) and minimal media (M9) |
| *Bacteroides thetaiotaomicron* | Gram-negative bacterium; present in mouse microbiome | Single condition |
| Matrices |  |  |
| Name | Type | Rationale |
| Mouse faecal pellets | High complexity | Most similar to real experimental conditions and challenges |
| Sterile corn roots | Low complexity | Low protein diversity; similar to situations observed, e.g., in symbioses |
| Gnotobiotic mouse faecal pellets | Medium complexity | Analysis of background complexity effects by comparison with complex mouse stool |

After proteins were extracted and digested, and peptide concentrations were determined as detailed in the Supplementary Methods, we mixed the resulting peptides of pure cultures in different known quantities with peptides extracted from one of the three matrices to create the final defined metaproteomes with known composition (see Results and Discussion Figure 1).

**Figure 1:**
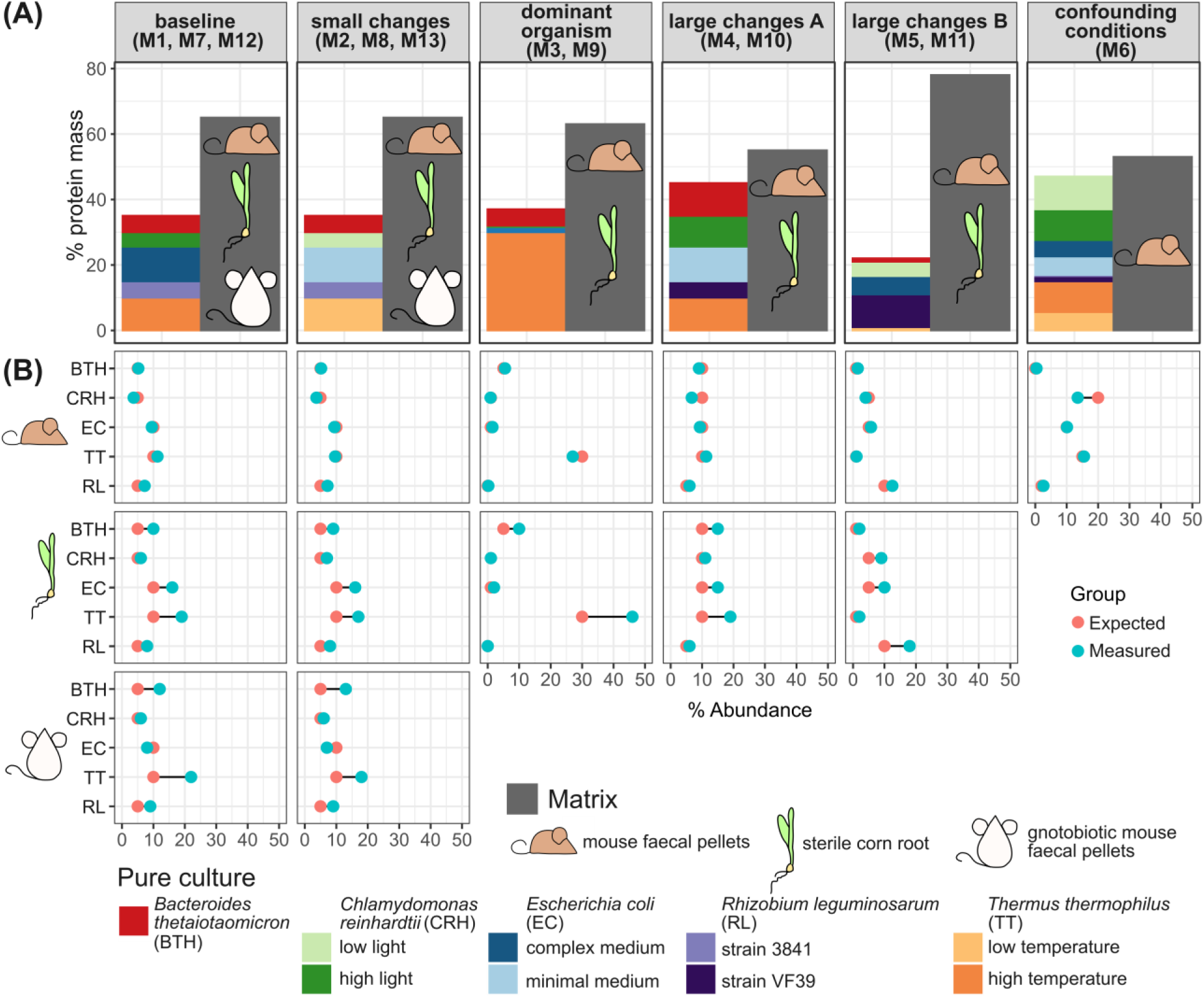
Design of defined metaproteomes and comparison of measured with pre-defined species abundances. All defined metaproteomes consisted of a metaproteome matrix with spiked-in pure cultures in known quantities. (A) Defined metaproteome compositions (see Supplementary Table 4 for percentage input per species). Grey bar corresponds to the peptide abundance of the respective matrix. We used one of three matrixes. For example, M1, M7, and M12 correspond to the same pure culture mixture added to either mouse faecal pellet protein (M1), sterile corn root protein (M7), or gnotobiotic mouse faecal pellet protein (M12). (B) Comparison of the measured species abundances (proportion of peptide signal summed per species to total signal) with their pre-defined input abundance (mass-% of peptides). Color-blindness accessibility note: the order of pure cultures in the legend corresponds to the order in the barplots (left to right columns = top to bottom).

### Metaproteomic database generation

The metaproteomic databases were assembled as a combination of “building blocks,” which included proteins expected to be found in the pure cultures, predicted mouse or maize proteomes, the matrix microbial translated metagenome, mouse dietary proteins (when applicable), and common laboratory contaminants (Supplementary Table 2). For the generic mouse stool matrix, DNA extraction and metagenomic analysis were performed (see Supplementary Methods) in order to generate a sample-specific database, as required for the metaproteomic analysis [32]. Each building block was clustered at 95% sequence identity using CD-HIT and then combined for the different mixes. Please refer to the Supplementary Methods for details.

### LC-MS/MS measurements and analyses

Peptides from the pure cultures and the different mixes were analyzed using a nanoLC-MS/MS system consisting of a Dionex UltiMate 3000 RSLCnano (Thermo Scientific) connected to an Orbitrap Exploris 480 (Thermo Scientific) equipped with an EASY-Spray source (Thermo Scientific). Samples were run in randomized block design. For each run, we loaded 1 µg of peptides per sample onto a trap column (Acclaim PepMap100, C18, 5 μm, 100 Å, 0.3 mm i.d. ×5 mm, ThermoScientific) and backflushed onto a 75 cm analytical column (EASY-Spray C18, 2 μm, 100 Å, 75 μm i.d., Thermo Scientific). Peptides were separated on the analytical column using a 140 min gradient of 95% eluent A [0.1% (v/v) formic acid], 5% eluent B [80% (v/v) acetonitrile, 0.1% (v/v) formic acid] to 31% (v/v) B in 102 min, 31 to 50% (v/v) B in 18 min, and finally to 99% B for 20 min at a flow rate of 300 nl/min. The mass spectrometer was operated in data-dependent mode and the 15 most intense peptide precursor ions were selected for fragmentation and MS/MS acquisition. The selected precursor ions were then excluded from repeated fragmentation for 25 s. The resolution was set to R = 60,000 and R = 15,000 for MS and MS/MS, respectively. As lock mass we used the ambient ion 445.12 m/z.

The obtained raw files were analyzed using their respective database (see Supplementary Methods), including decoy sequences, with the Sequest HT node in Proteome Discoverer version 2.3.0.523. We set peptide mass tolerance to 0.1 Da and fragment mass tolerance to 10 ppm. Trypsin was set as digestion enzyme, and two missed cleavages were allowed. As fixed modification, we set carbamidomethylation of cysteine residues, and as variable modification, protein N-terminal acetylation, oxidation of methionines, and deamidation of asparagine and glutamine. We filtered identifications for a protein false discovery rate (FDR) of 5%. The outputs of samples belonging to one matrix (all replicates and mixes generated with that matrix) were reprocessed using the consensus workflow of Proteome Discoverer, leading to one protein group file per matrix.

### Detection of differential abundances

We filtered, imputed, then normalized and transformed metaproteomics quantification data, and subsequently applied tests for statistical inferences as summarized in Table 2. Please refer to the Supplementary Methods for details on data preprocessing, filtering, imputation, normalization and transformation, and statistical tests used. We used R 4.3.1 [33]. For details on package versions, please refer to Supplementary Table 3.

**Table 2:**
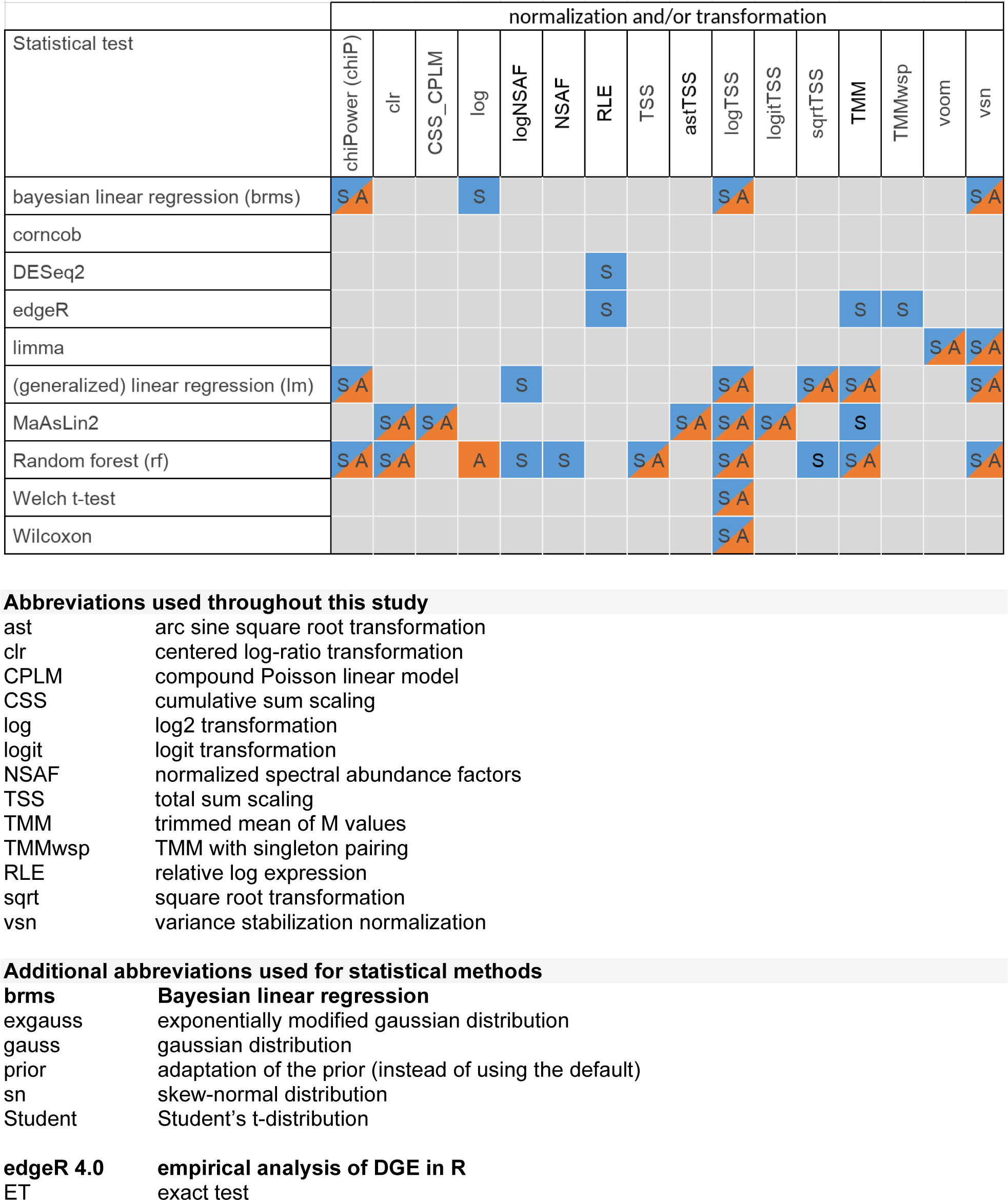
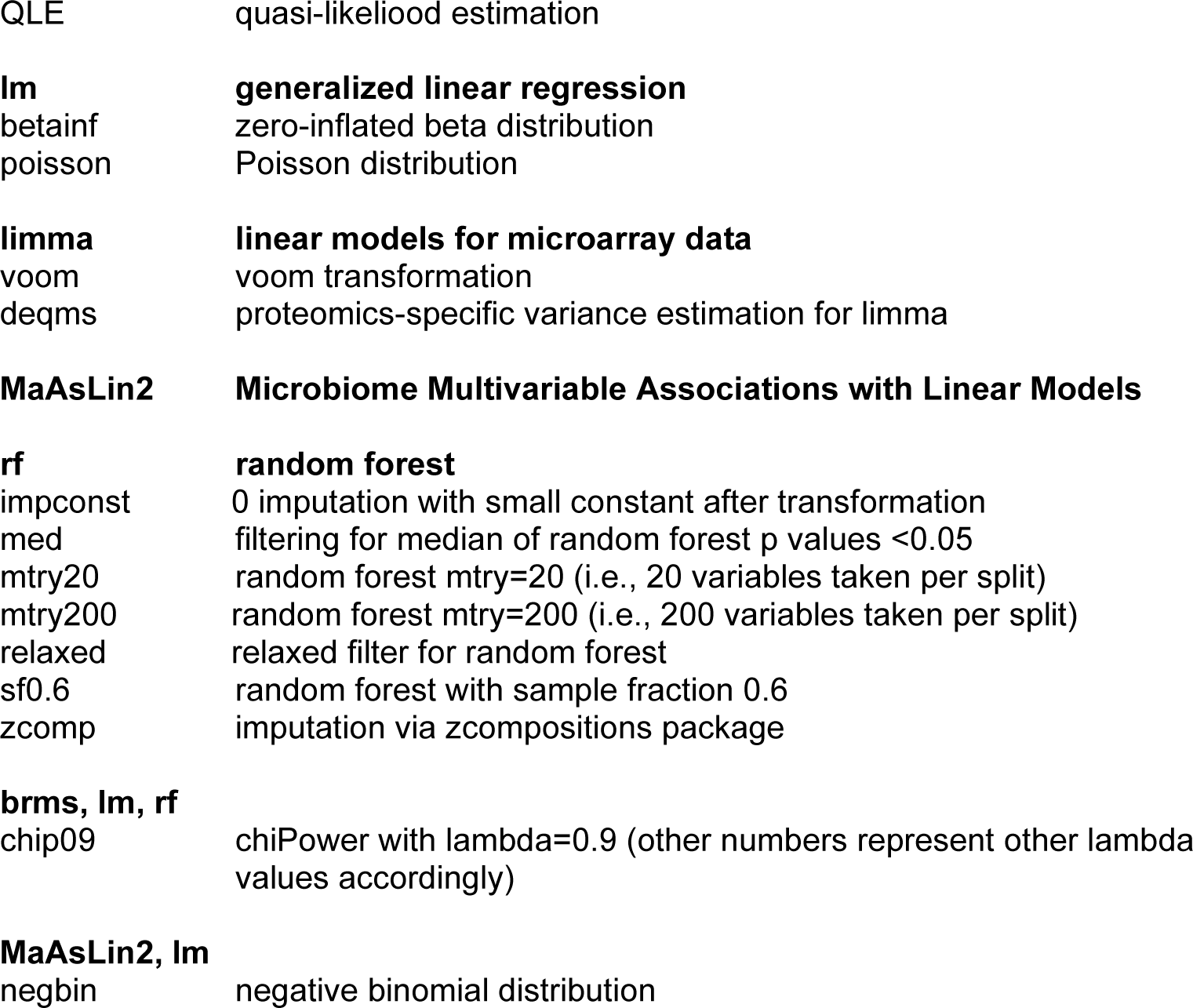
Combinations of statistical tests and normalization (+transformation) methods used in this study. Note that only certain normalization/transformation and test combinations are compatible, as given below. The method corncob does not normalize beforehand. S: used for spectral count data, A: used for AUC data. Abbreviations correspond to abbreviations used throughout the study, please see below table. For details and references, please refer to the Methods section.

## Results and Discussion

### Design of defined metaproteomes and statistics evaluation framework

We designed defined metaproteomes with varying degrees of complexity to evaluate differential expression analysis statistical tools and methods. We built these defined metaproteomes by mixing defined amounts of peptides extracted from pure cultures with peptides extracted from one of three complex matrices. The three matrices included faecal material from conventional mice representing a high-complexity microbiome with hundreds of species, faecal material from gnotobiotic mice with a defined 13-species microbiome representing a medium-complexity microbiome, and sterile corn roots representing a low-complexity microbiome after mixing in the pure cultures. For the pure cultures, we grew five microbial species, namely *Bacteroides thetaiotaomicron*, *Chlamydomonas reinhardtii*, *Escherichia coli*, *Rhizobium leguminosarum*, and *Thermus thermophilus* in one or two different culturing conditions, or used different strains of the same species (see Results and Discussion Figure 1). For the species grown in two conditions we chose conditions that induce distinct protein abundance profiles. The resulting defined metaproteomes corresponded well between the actual measured species abundance and the theoretical (pre-defined) input abundances for each species. This shows that the sample generation process was successful and that these samples can be used to determine the performances of statistical tests considering various metaproteomic data challenges (Figure 1).

For testing statistical data analysis methods, we used regression-based approaches, machine-learning based techniques, and null hypothesis testing (Table 2). Different tests are compatible with different normalization and/or transformation techniques (Table 2 and Methods). When applicable, we set the nominal false discovery rate (FDR) for detection of significant differences to 5 %. The FDR measures how many of the detected significantly different variables are, in fact, not significantly different between conditions.

We based evaluation of statistical tests for the defined metaproteomes on metrics derived from true positive (TP), true negative (TN), false positive (FP), and false negative (FN) identifications (Figure 2). The ground truth used for these calculations depended on the type of comparison:

i. If normalizing for species abundance and comparing between different culturing conditions of an organism, we defined as ground-truth those proteins that were statistically significantly different between the culturing conditions when measuring the pure cultures. Proteins that were only detected in pure cultures, but not in the defined metaproteomes, were not taken into account. Few protein groups were only detected in defined metaproteomes, but not in pure cultures. We removed these from our analyses, as we were not able to assess based on our definition whether these are true or false positives/negatives. This comparison applies, for example, for *T. thermophilus* between M2 and M3 (Figure 1).
ii. If not normalizing for species abundance, but comparing proteins of one species between conditions, with the same total species abundance in both mixes (i.e., no shifts in relative protein abundances which are solely caused by the different relative abundances of the organism), the ground-truth was comprised of the same proteins as in (i). One example is the comparison of *T. thermophilus* between M1 and M2 (Figure 1).
iii. If normalizing for species abundance and comparing proteins of one species within the same condition, or not normalizing and comparing between the same condition and same abundance, there are by definition no TP, and all proteins that are detected as significantly different are by definition FP. See, for example, *T. thermophilus* M1 vs. M3, or *T. thermophilus* M1 vs. M4 (Figure 1).
iv. If not normalizing for species abundance for a specific species grown under one condition, and the abundance of that species was different between mixes, then by definition all proteins from that species are differentially abundant and can be considered TP if detected as statistically significantly different. In this comparison, there are no FP or TN, but only FN, i.e., those proteins of the species which were not detected as being significantly differentially abundant. This type of comparison occurs, for example, for *T. thermophilus* M1 vs. M3 (Figure 1).
v. The comparison of no normalization for species abundance, but changes in condition and species abundance cannot be evaluated, because it is not known whether the changes are due to species abundance changes, condition changes, or both (i.e., changes in abundance of the organism are confounded with changes abundance of individual proteins due to different culturing conditions). We assigned “not available” (NA) to these values. Please note that in a real-life dataset, where the abundances of organisms are usually not known beforehand, it is thus necessary to normalize on organism (or sub-community) level, if biologically meaningful conclusions on the organism level should be drawn (see also Kleiner, 2017). This pertains, e.g., to *T. thermophilus* M2 vs. M3 (Figure 1).
vi. We also assigned “NA” to TP and TN where the organism was not present in both conditions, as all detected proteins are already wrongly detected; there is no ground truth for this type of comparison. See, for example, *R. leguminosarum* strain VF39 in M1 vs. M2 (Figure 1).

**Figure 2:**
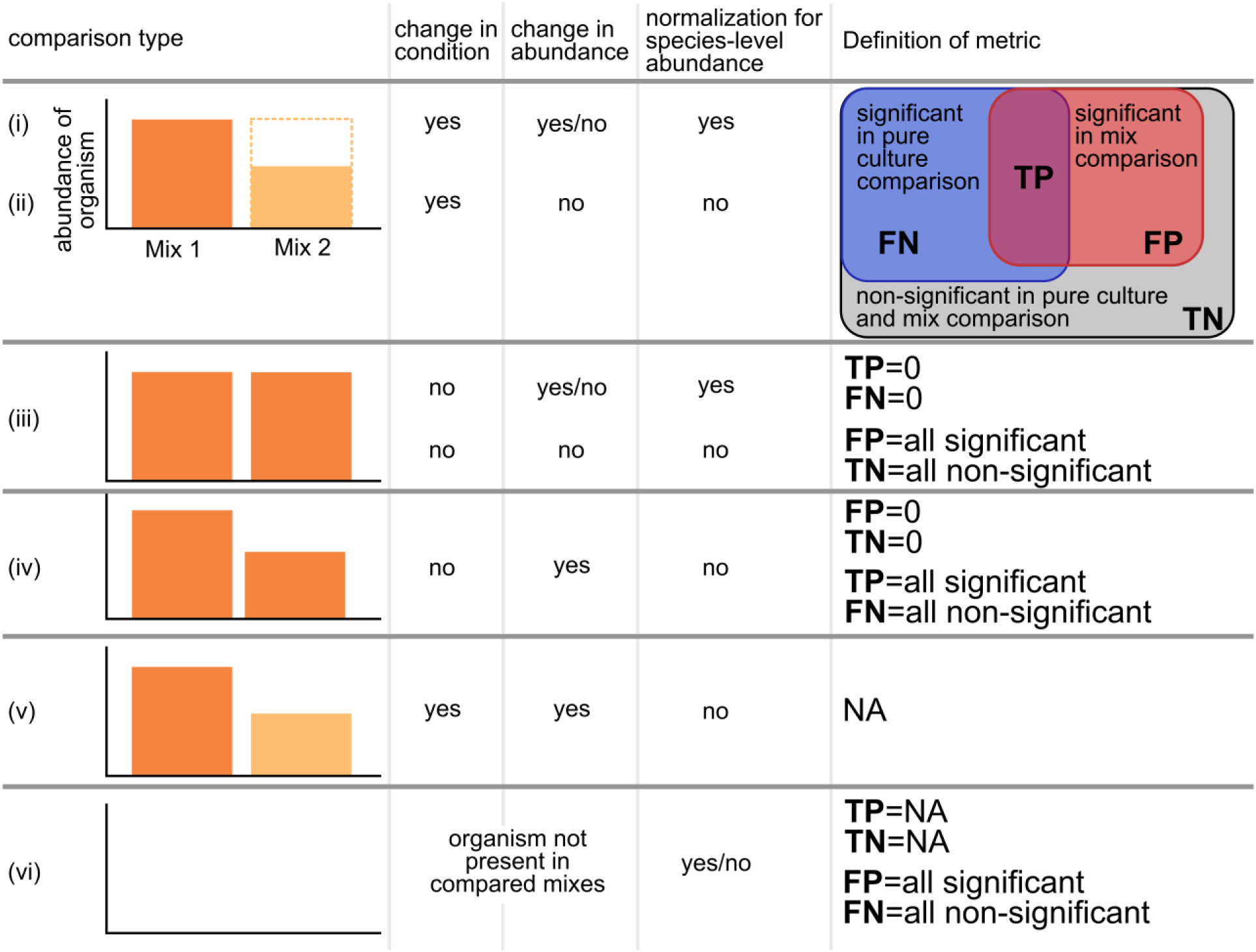
Definition of outcomes for each metric used for performance assessment of statistical tests in this study. For reference to scenarios, please refer to the text. Dark vs. light orange: Culturing condition 1 vs. culturing condition 2.

For comparison types (i) and (ii), the final evaluation was based on the following metrics (all ranging between 0 and 1, and chosen so that they will be 1 for ideal conditions):

1. Positive predictive value (PPV) = Precision = complement of the false discovery rate (FDR; [34, 35]):

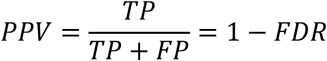
2. Negative predictive value (NPV) = complement of the false omission rate (FOR):

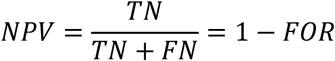
3. True positive rate (TPR) = sensitivity = recall = complement of the false negative rate (FNR):

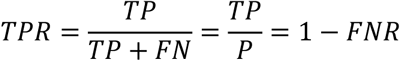
4. True negative rate (TNR) = specificity = complement of the false positive rate (FPR):

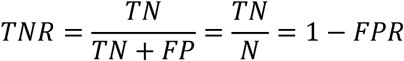
5. Balanced accuracy (BAcc; [36]):

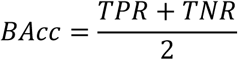
6. F1 score = harmonic mean of precision and recall [37]:

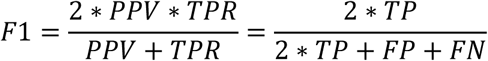
7. P4 score [38], which balances the PPV, TPR, TNR, and NPV (i.e., if the P4 score is close to one, all of the metrices used are close to one):

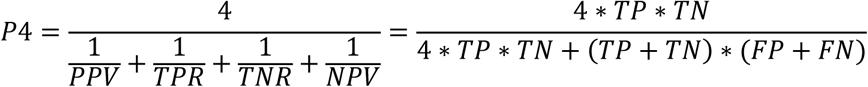

For the comparisons (iii) to (vi), calculation of these metrics does not make sense, as some (or all) of the base metrics (TP, FP, TN, FN) are 0 or NA.

### Determining statistical tests for pure culture ground truth differentially abundant proteins

We used pure culture proteome data to identify protein groups whose differential abundances between mixes were sufficiently reliable to use for downstream analyses of statistical test performance. For that, we determined the ground truth of statistically significantly different protein groups between conditions using the proteomes of the pure cultures that were the inputs for the defined metaproteomes. To do this, we first determined which statistical test to use for pure culture proteome comparisons. To determine this test, we used the defined metaproteomes with the mouse fecal pellet matrix (M1 to M5) that had EC, TT, and CRH at the same condition, but at different abundances, meaning that for each of these three species all proteins should be statistically significantly different in the respective tests, when not normalizing on the species level. We then identified the tests yielding the best balance between true positives and true negatives, by increasing thresholds for TN and TP until one test remained. For SpC, this led to a threshold of 70% TP and 95% TN, resulting in us using brms_gauss_chiP09 (Figure 3) to identify differentially abundant protein groups in pure cultures to be used as ground truth (Supplementary Tables 5a, 5b). For AUC, we used thresholds of 70% TP and 85% TN. We chose lm_vsn as pure culture ground-truth test, because while both lm_vsn and brms_gauss_vsn gave the same performance, lm_vsn is computationally less intensive (Supplementary Tables 6a, 6b). To ensure that our results and conclusions are not biased due to our choice of ground truth test, we determined the overlap between significant proteins detected with several well-performing tests (Supplementary Figure 6 and Supplementary Results and Discussion) and the effect of a changed ground truth on performance in metaproteomes (Supplementary Figure 7 and Supplementary Results and Discussion). We found that there was large overlap in proteins detected as significant between all tests particularly for AUC data, and that changing the ground truth tests only led to minor changes in the ranking of tests that performed well on metaproteomic data.

**Figure 3:**
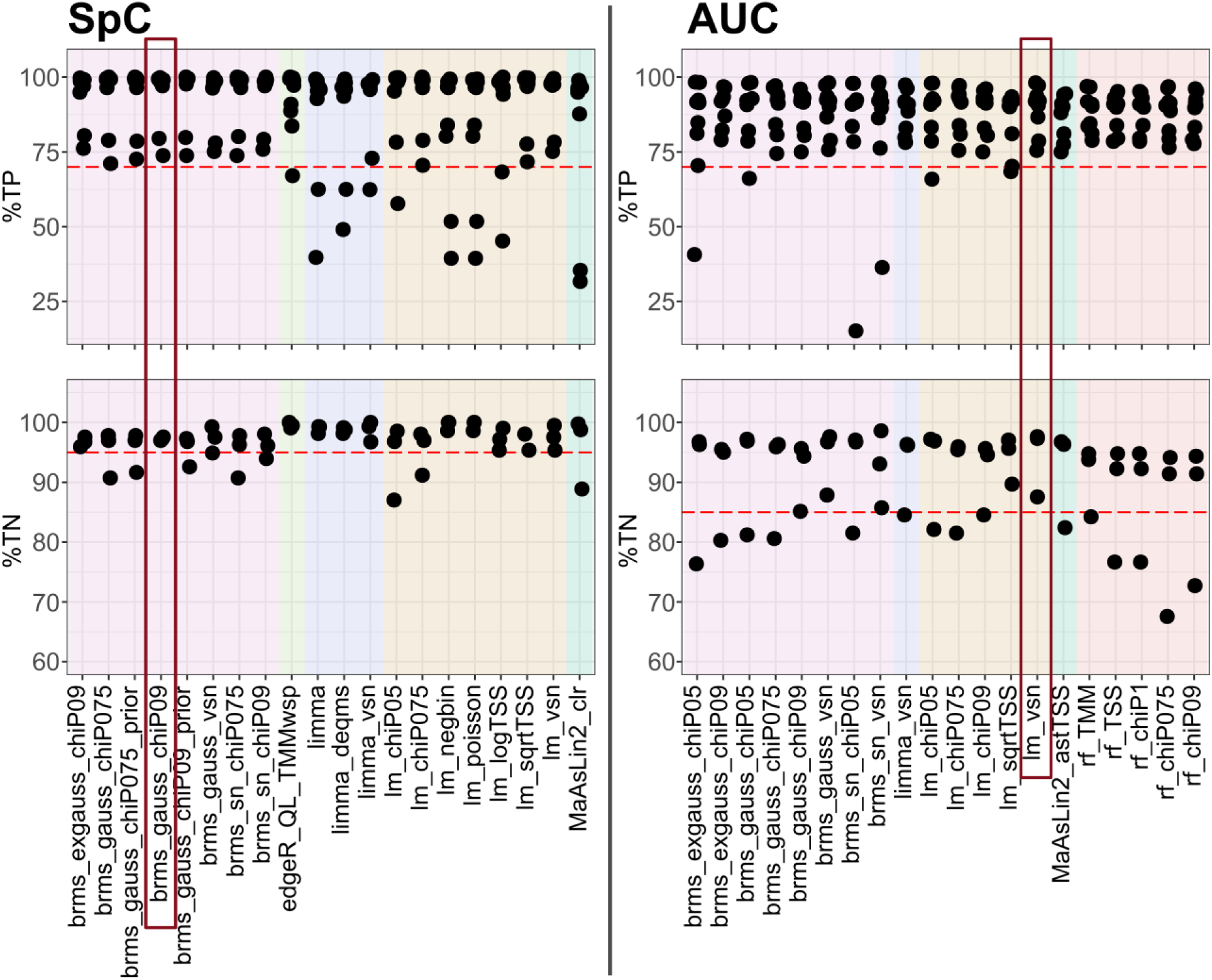
Determination of the ground-truth test for evaluating statistically significantly different proteins in pure cultures. We pre-selected tests with a median %TP and %TN of 95 for SpC, and of 90 for AUC data. %TP and %TN: Percentage of true positives/ true negatives of all proteins of the respective organism (T. thermophilus, E. coli, or C. reinhardtii) in comparisons where the expectation was that all (100%) proteins of an organism would be differentially abundant, because the organism was mixed into the compared samples at different abundances (but same condition). Red boxes indicate the tests used to identify differentially abundant proteins in the pure cultures, which we define as the statistical ground-truths. Red dashed lines: 70% TP and 95% TN for SpC, 70% TP and 85%TN for AUC, respectively.

### Many statistical approaches perform similarly well, and some perform poorly

With the determined ground truth of statistically significant proteins, we next determined how each statistical test was able to identify differentially abundant proteins in the mouse faecal pellet mixes 1 to 5, using the results of comparison types (i) and (ii) as outlined in Figure 2. No single statistical approach clearly out-performed the others (Figure 4). Please note that the scores we chose can range from 0 to 1 and trend towards 1 for desirable performance outcomes. Some statistics clearly performed worse than the majority of tests (e.g., Wilcoxons rank-sum test, DESeq2) as well as some normalization methods tested for random forest implementations. Based on these results we narrowed the list of tests to use in further evaluations to well-performing tests, which included tests with a median TNR and TPR in the top 25% of all tests. Since some tests with high median performance suffered from a large spread in TNR and TPR, we added the requirement that the the TNR and TPR lower quartiles for each test had to be greater than those of 25% of all tests (Supplementary Tables 5c, 5d, 6c, 6d). This resulted in us retaining 30 tests for SpC-based approaches (42% of all 72 tests), and 17 of 47 (36%) for AUC-based tests (Figure 4). Due to the fixed cutoff, some tests were removed, which performed only slightly less good than these selected tests. Generally, we noticed that the NPV performed worse than all other metrics, meaning that all tests miss a large share of actually significant differences (Figure 4). Our approach was largely robust to changes in the ground truth test (see Supplementary Results).

**Figure 4:**
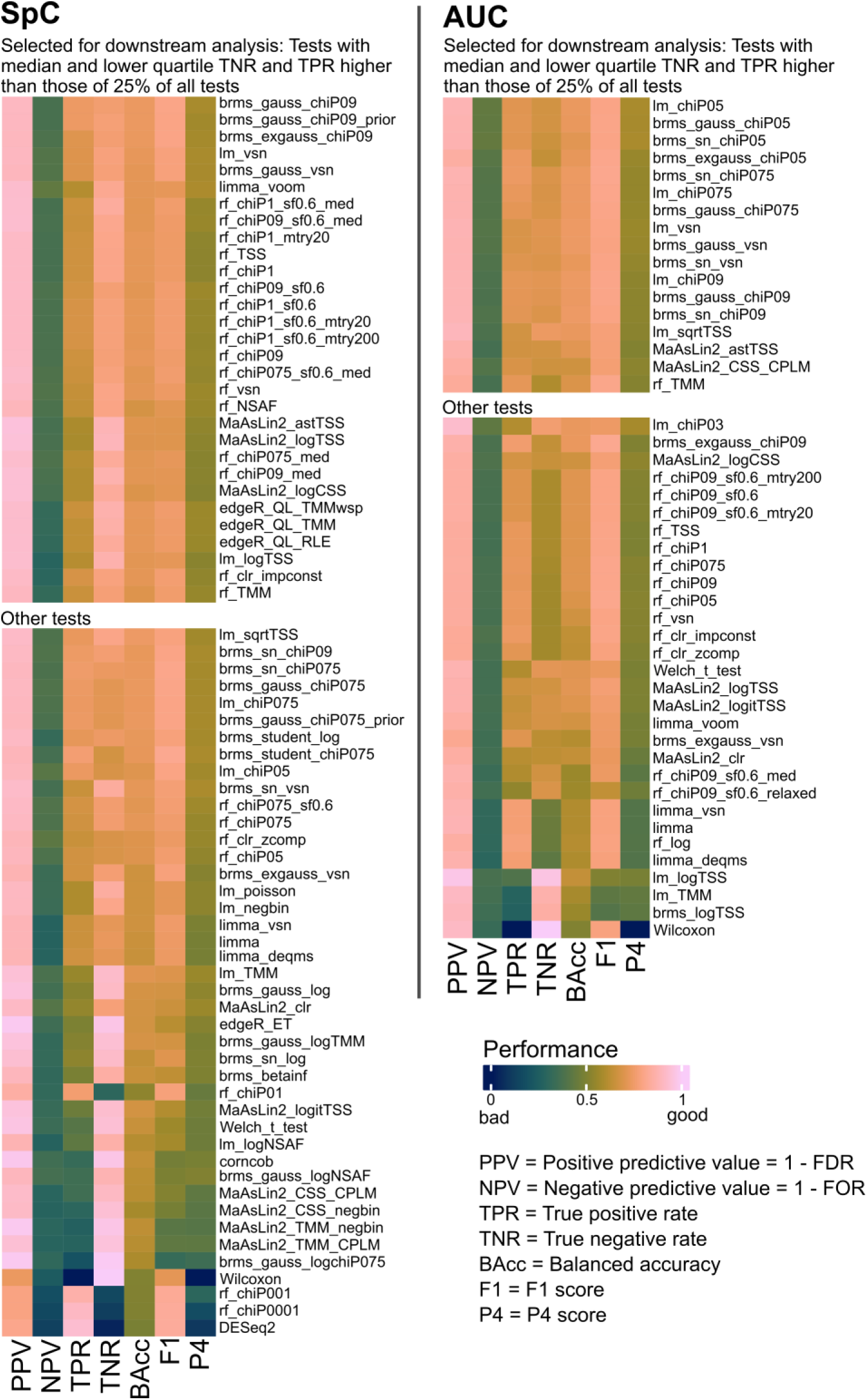
Median test evaluation metrics for mouse faecal pellet matrix comparisons (M1 to M5). Upper part: tests with a median and lower quartile TNR and TPR higher than those of 25% of all tests. The tests in the upper part were used for further in-depth analysis. Data was clustered row-wise based on Canberra distances.

Our results partially corroborate comparisons of statistical approaches that were done on other (meta)-omics data types. We found that compositional analyses methods generally did not perform better than methods not explicitly taking compositionality into account. However, methods that take compositionality into account, for example, chiPower transformation, were also among the best performing methods. Thus, based on our data, explicitly accounting for compositionality is not a requirement for all combinations of statistical tests and data preparation even though it can improve results for some tests. This is in line with previous results from a study comparing statistical approaches for 16S rRNA gene amplicon and WGS data [19]. However, in that study, DESeq2, limma-voom, and corncob were recommended as best-performing, while in our study DESeq2 and corncob were among the worst-performing approaches. Another study on use of statistical approaches for RNA-Seq data found that DESeq2, limma-voom and edgeR (version 3) produced too-high false positive rates and the Wilcoxon rank-sum test was recommended [39]. In contrast, we found that limma-voom performed well on our data, and Wilcoxon rank-sum test performed poorly. These contrasting results between studies comparing statistical approaches for omics data are likely due to several causes, including 1) differences in underlying data structures of different (meta)-omics data types despite some similarities (e.g., compositionality and sparsity), and 2) different frameworks for evaluating performance of statistical approaches. While some past studies focused heavily on FPRs or FDRs (=1-PPV) (e.g., [19, 39], we used a balanced framework: we took into account not only true and false positives, but also true and false negatives, i.e., PPV and NPV (as complements of the FDR and FOR, chosen so that they will approach 1 for a desired outcome), as well as TPR and TNR. For final scoring, we compared the F1 score to the more balanced P4 score, and focused on the latter. For example, if we focused only on TNRs, the Wilcoxon test would be a clear winner, however, when also considering TPRs, it becomes immediately clear that this test is unsuitable. The same principle holds true for DESeq2, which fares well with regard to the TPR, but had the lowest TNR. The tests we chose for more in-depth analyses show a more balanced performance, and thus decrease the risk for a biased data analysis.

### Type of comparison and data pre-treatment impact test performance

For the 30 SpC tests and 17 AUC tests for which we carried out in-depth analyses, we found that the type of comparison, in terms of organisms, as well as in terms of organism abundance compared, had a major impact on performance, as demonstrated by the large spread of P4 scores for individual tests (Figure 5, Supplementary Figure 2 for all matrices). The type of mix impacted the outcome more than the statistical test chosen: the intraclass correlation (ICC), reflecting the similarity of test results, was (for SpC) 0.89 (F-test, p < 0.01) between tests, but only 0.02 (F-test, p < 0.01) between mixes. For AUC, the ICC between tests was even higher at 0.95 (F-test, p < 0.01) and lower between mixes (0.006, F-test, p < 0.01).

**Figure 5:**
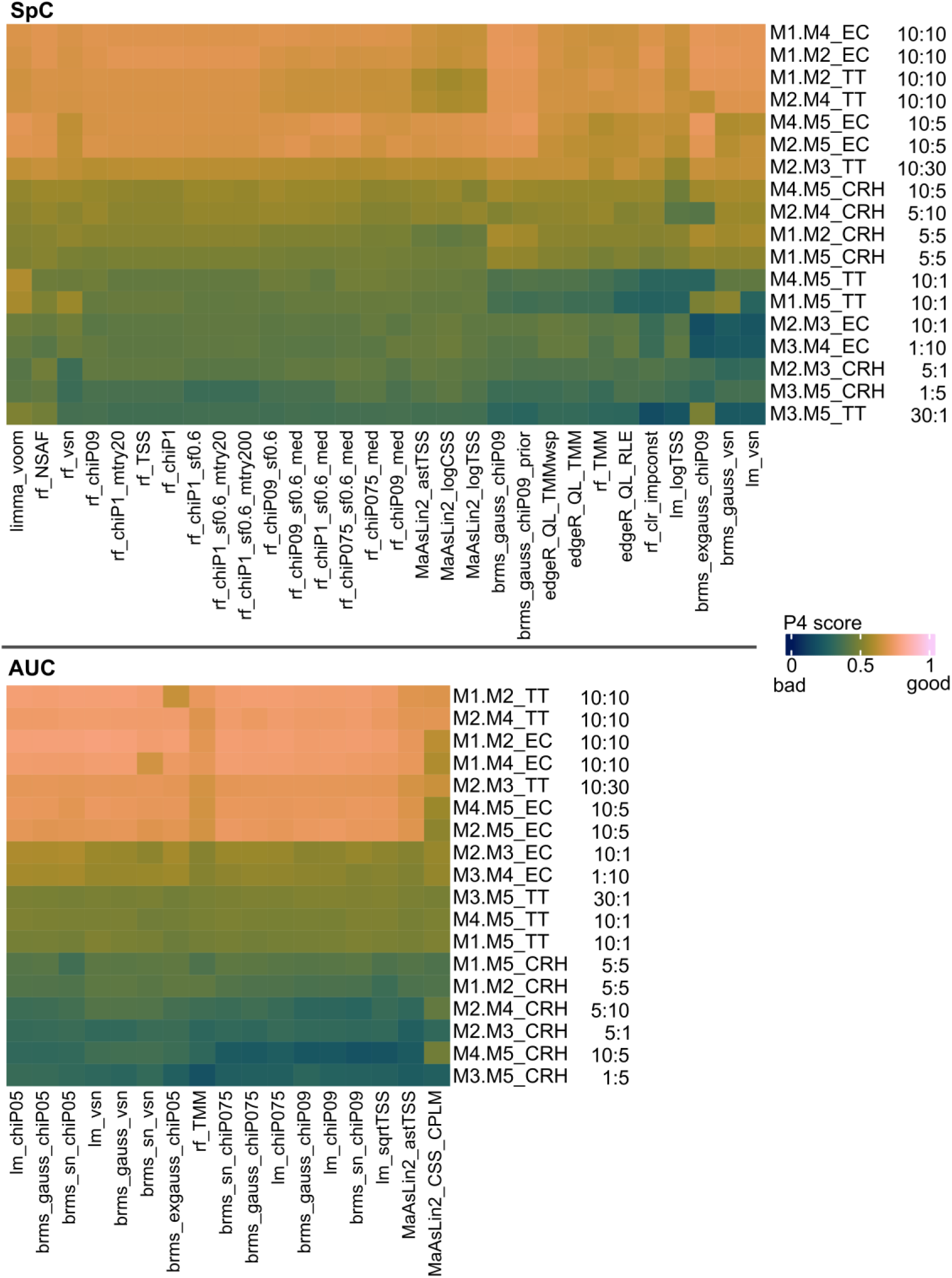
Heatmaps of P4 scores for the best-performing statistical approaches in mouse faecal pellet matrix (comparisons M1 to M5). Tests shown here were chosen as the best-performing ones based on data shown in Figure 4. The P4 score ranges from 0 to 1 and trends towards 1 for desirable performance outcomes. Data was clustered row- and column-wise based on Canberra distances. CRH: Chlamydomonas reinhardtii, EC: Escherichia coli, TT: Thermus thermophilus. Numbers to the right give the abundance of the respective organism in percent in the compared mixes.

Specific comparison types that led to low performance, indicated by low P4 scores across tests, involved species that were present in the defined metaproteome samples at 1%. We observed this low performance for low-abundance species for both SpC and AUC-based analyses. This low performance of tests on samples with low-abundance species is likely due to more inaccurate quantification during LC-MS/MS analysis. Additionally, all comparisons that involved *C. reinhardtii* had low P4 scores, especially for AUC analysis, as compared to comparisons including *T. thermophilus* and *E. coli*, even when considering comparisons with the same species abundance ratios (see Supplementary Figure 2 and Supplementary Tables 5 and 6 for all matrices). Notably, *C. reinhardtii* showed very low numbers of proteins constantly identified across all mixes in the SpC data compared to its overall high number of proteins identified. This lack of overlap likely explains the low P4, caused by a low NPV (Supplementary Tables 5c, 5d, 6c, 6d and Supplementary Figure 3). For the AUC data, on the other hand, overlap between identifications was very high (Supplementary Figure 4), indicating protein mis-identifications and thus AUC quantification distortions, which might have been caused by the transfer of precursor peptide identifications between datasets (i.e., match between runs), as performed in ProteomeDiscoverer (see manual), but also in other software such as MaxQuant [40].

While median test performances and P4 scores mainly clustered by the statistical test itself (i.e., most random forest implementations clustered together, as did most limma, etc.), normalization and transformation did in some cases have a noticable impact as well (Figure 4, Figure 5): the voom normalization greatly increased performance of limma, while chiPower pretreatment with a too-low lambda worsened the random forest performances. On the other hand, random forest performance was at most slightly impacted by across-forest summarization via kernel density vs. median statistics or hyperparameter tuning. For edgeR, a quasi-likelihood estimation outperformed the exact test statistics (Figure 4).

Explicitly accounting for compositionality, e.g., via a clr transformation, has been advocated as mandatory for microbiome datasets [27]. According to our analysis, normalization and transformation impacted test performance, but accounting for compositionality was not necessary and in fact use of clr transformed data (but not of chiP transformed data) led to worse performance (Figure 4). Similar effects were shown by a study on amplicon sequencing data, where clr, isometric log-ratio, and additive log-ratio transformations performed slightly worse as compared to normalizations which do not take compositionality into account [41].

For random forests, imputation with a small constant performed slightly better than imputation based on compositionality (zcomp), indicating that more intricate methods are not always necessary (Figure 4). While more complex or data-dependent imputation methods exist (see, e.g., Lazar et al. 2016; Webb-Robertson et al. 2015), as well as methods explicitly taking into account the pattern of missingness, we found these to be unsuitable for the low number of replicates often available in metaproteomic studies and we are therefore not addressing these here. More stringent pre-filtering led to more true positives, but had inconsistent effects on the number of true negatives (see Supplementary Results and Supplementary Figure 1).

### Best tests per test category highlight test-specific differences

To determine the set of best-performing tests for more detailed analyses, we subsetted the tests to only include the test with the highest median PPV (=lowest FDR) per test category (i.e., brms, edgeR, limma, lm, MaAslin2, and random forests; edgeR and limma are only applicable to SpC), in the well-performing tests for all comparisons involving M1 to M5. This resulted, for SpC, in the six tests brms_gauss_chiP09, edgeR_QL_RLE, limma_voom, lm_logTSS, MaAsLin2_astTSS, and rf_chiP09_med. For AUC, we included the four tests brms_sn_chiP09, lm_sqrtTSS, MaAsLin2_astTSS, and rf_TMM (Figure 6, Supplementary Tables 5e, 6e). The spread of results per test is still wide, underlining that the specific comparisons performed impact the results. Additionally, the F1 score showed more differences between tests as compared to the P4 score, highlighting that the F1 score is only impacted by the PPV and TPR, whereas the P4 score is a balanced score of PPV, NPV, TPR, and TNR. The general result of high PPV but rather low NPV, as discussed above, holds true here as well. For SpC-based tests, brms_gauss_chiP09 clearly outperformed the others in terms of the TPR, but had a slightly lower TNR. On the other hand, limma_voom had the highest median NPV, and the lowest performance spread of the TNR. The AUC-based tests generally had a lower median PPV and larger spread of PPV (and most other metrics) compared to the SpC-based tests. Within the AUC-based test, differences between medians were mostly lower than for SpC-based tests. The performance of all tests decreased for comparisons with confounded data (i.e., with M6) as expected (Supplementary Results and Discussion, Supplementary Figure 5, Supplementary Tables 5f, 6f).

**Figure 6:**
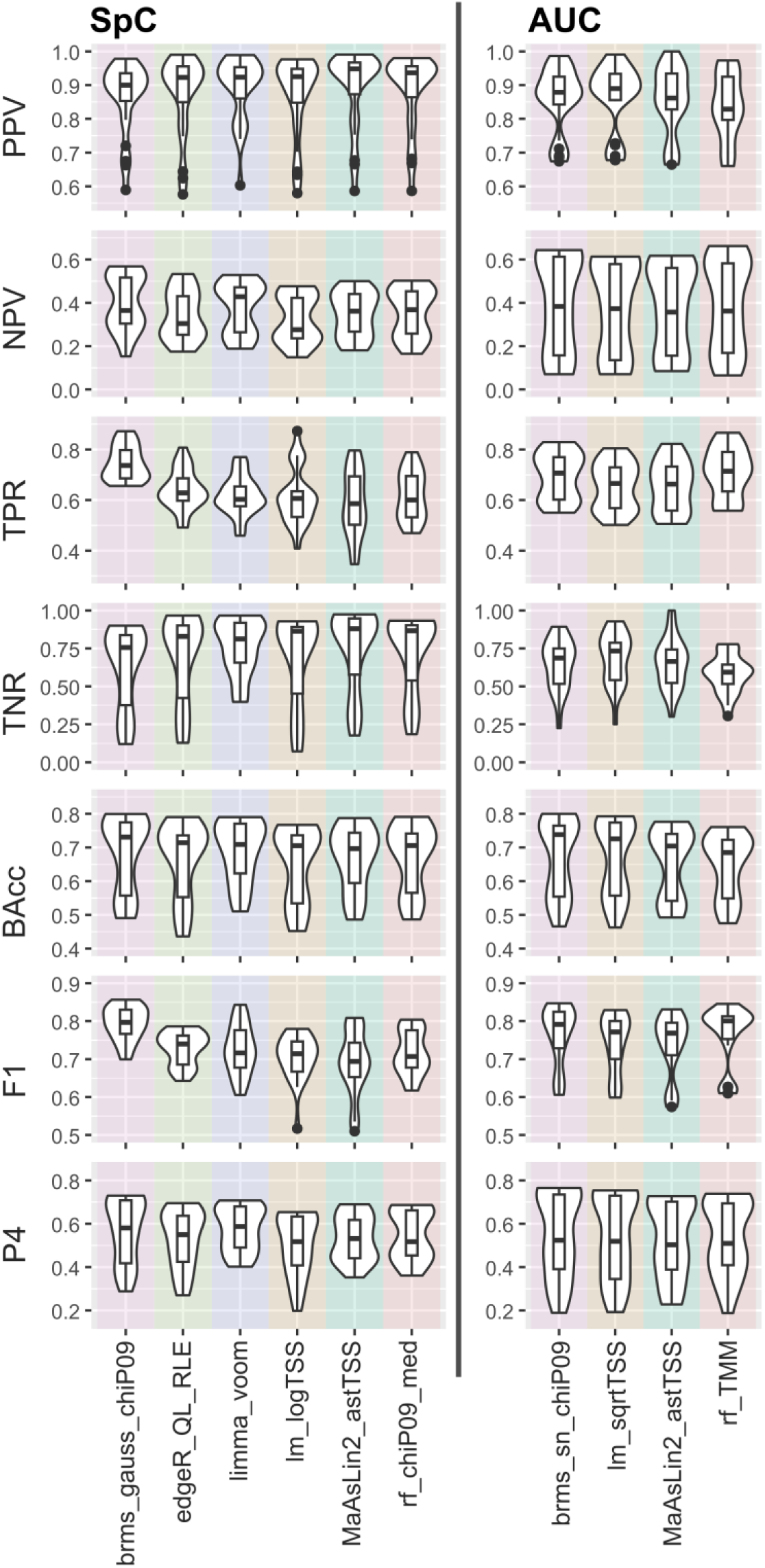
Test metrics for the best-performing tests in the mouse faecal pellet matrix defined by metaproteomes (M1 to M5). Shown are median values (boxplots) and value distributions (violins).

### Protein (mis)identification effects

One challenge in metaproteomics is that proteins from closely related strains/species are hard to differentiate as large portions of their protein sequences are identical, which makes it less likely that differentiating peptides with differences in amino acid sequence are identified [2]. This can lead to misidentifications where the correct protein is identified, but it is assigned to the wrong species/strain, which can confound statistical tests. To address misidentifications between different bacterial strains, as well as between pure culture and matrix microbiome, we used the two *Rhizobium* strains RLVF39 and RL3841, and *B. thetaiotaomicron*, a microbial species that is also present in the mouse stool naturally (Figure 7). These analyses included comparison types (iii) to (vi) as outlined in Figure 2. Comparisons involving a higher protein abundance of one organism increased test performance for RLVF39 and RL3841 comparisons. Considering the TN, organism-level normalization generally (slightly) increased test performance (as expected). Performance decreased visibly for SpC data using the lm_logTSS considering RL3841 in the M1 vs M2 comparison (i.e., comparing 5% abundance each), likely because the model fit was comparatively low (a test of taking all *Rhizobium* proteins in this comparison together, i.e., including potential strain misidentification, produced a boundary fit, i.e., the random effect was estimated as zero).

**Figure 7:**
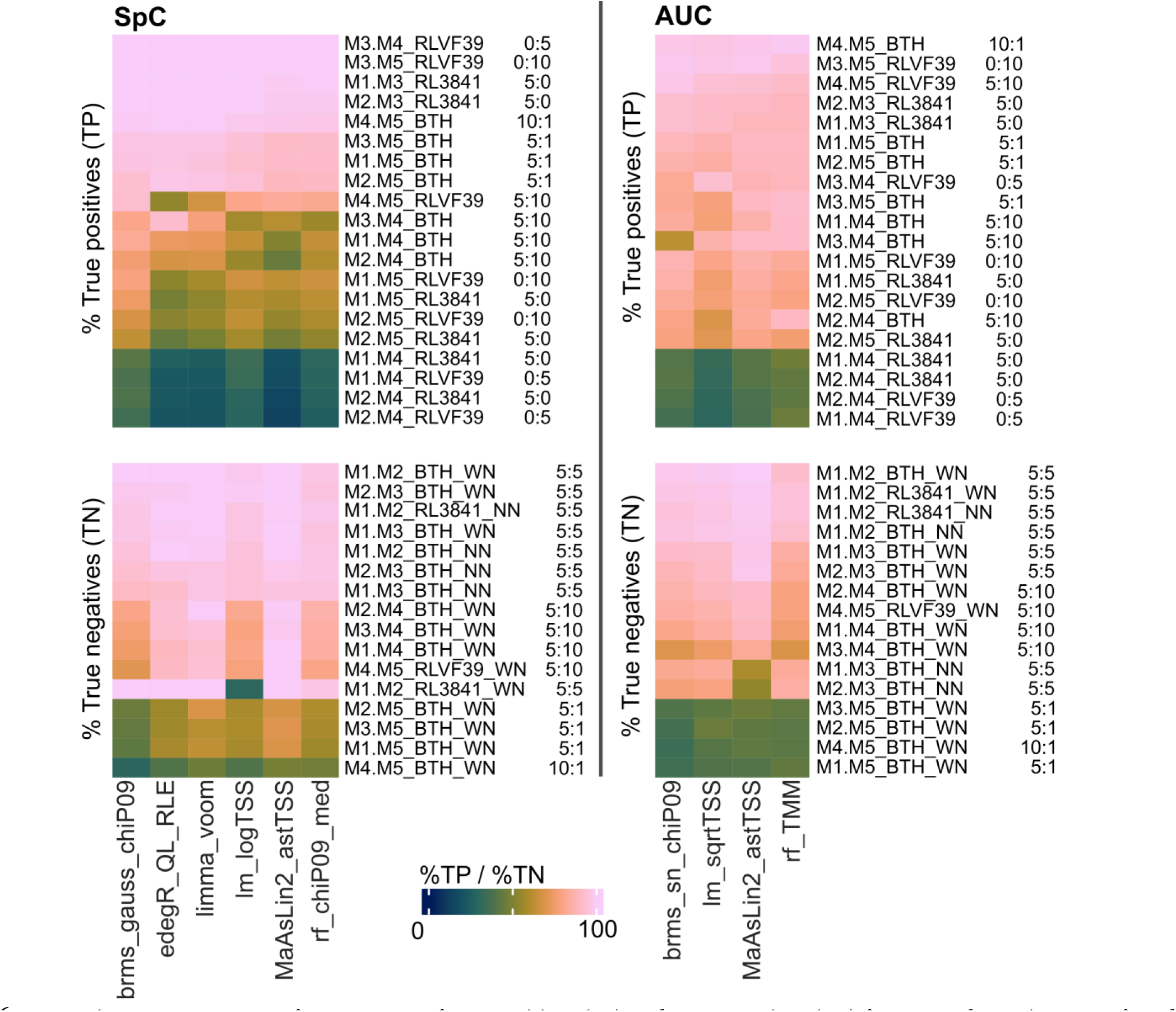
Heatmaps of percentages of true positives (TP) and true negatives (TN) for pure cultures in mouse faecal pellet matrix (M1 to M5) without different culturing conditions (B. thetaiotaomicron and the two Rhizobium strains RL3841 and RLVF39). Numbers give the abundances of the respective organism in percent in the compared mixes. Data was clustered row- and column-wise based on Canberra distances. WN: with organism-level normalization, NN: no normalization on organism level. Note that no normalization is only possible for evaluation of TN; for TP to be present, there has to be an actual abundance change, and when normalizing, this change in abundance is by definition eliminated.

### Matrix complexity effects

The three matrices we used for the defined metaproteomes mainly impacted the NPV and, concomitantly, the TNR (Figure 8). While performances of the best tests were similar for SpC across all three matrices, performances of the best AUC tests varied widely between matrices. The gnotobiotic mouse matrix generally provided best performance - but it has to be noted that this included only one type of comparison, i.e., between M12 and M13, thus excluding many of the challenges present for comparisons within the other two matrices. Based on the P4 values per comparison and test (Supplementary Figure 2, Supplementary Tables 5e, 5h, 5i, 6e, 6h, 6i), comparisons involving CRH and very low-abundant organisms (i.e., 1%) were especially challenging for AUC based tests, independent of the matrix.

**Figure 8:**
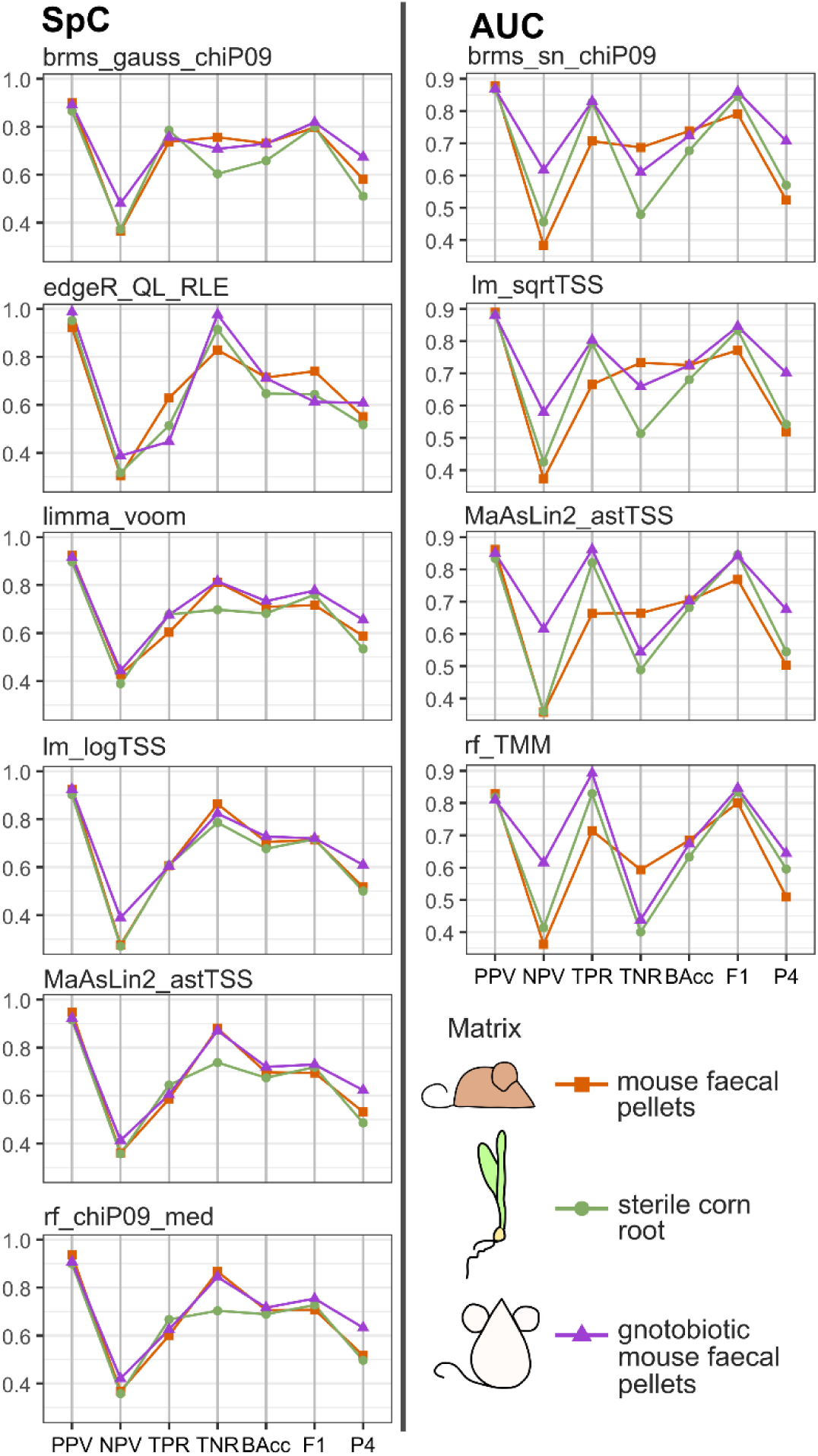
Evaluation metrics for best tests for all three matrices used. Shown are parallel coordinates plots with median values (please note that, while the mouse faecal pellet matrix and sterile corn root matrix include the same number and kind of comparisons, in the gnotobiotic mouse faecal pellet matrix only M12 and M13 are present limiting the number of comparisons to one). Detailed P4 score comparisons for data from corn root and gnotobiotic mouse matrix are shown in Supplementary Figure 2.

## Conclusion

In this study, we developed a complex and controlled sample set and performance test framework to evaluate statistical approaches for differential abundance analysis in metaproteomic studies. Our known ground truth enabled a “gold standard”-based evaluation of statistical approaches for data-dependent metaproteomics with n=4 replicates, a common replication level in environmental metaproteomics studies.

Pertaining to the general challenges for metaproteomics data analysis we designed our samples for, tests generally fared worse at (a) detecting small changes, than (b) detecting large changes without increasing false discoveries. The challenge (c) of misassignments becomes relevant especially for related strains, and, while thus pertaining to all tests, was apparently somewhat better controlled by brms_gauss_chiP09 for SpC data, whereas for AUC, there was no clear difference between the best-performing tests.

Regarding differences between tests, SpC-based tests fared better than AUC-based tests at controlling the PPV (=1-FDR), i.e., detecting proteins as significantly differentially abundant which are truly differentially abundant. The NPV (=1-FOR) was low for all tests which passed initial quality assessment criteria, which means that many protein groups that were in fact significantly differentially abundant were not detected as significantly different. While in many studies significant differences between conditions are the focus of analysis, and will be interpreted in greater detail, this low NPV (=high FOR) means that we are missing a lot of true differences as they do not show up as significant in the tests. This certainly needs to be taken into account when drawing conclusions from metaproteomics data. A low NPV is, however, not a metaproteomics-specific issue, as it is likely low in many (meta-)omics approaches. NPV, however, is often not evaluated for omics-focused statistics due to the absence of a known ground truth for true positives. Our results highlight that evaluations of statistical approaches for omics data should consider a framework that enables the assessment of NPV/FOR.

One way to increase the NPV is to use a higher number of replicates, which is often difficult in metaproteomics studies that use limited environmental samples or samples from small cohorts (e.g., prospective clinical studies). For example, RNA-Seq needed at least 20 replicates per condition to identify 85% of the truly differentially expressed genes, and it has been recommended to use at least 6 replicates per condition [43]. Increasing biological replication should thus always be considered when feasible. Additionally, the impact of number of replicates on test performance is a factor worth studying further for metaproteomics.

The performance of AUC-based tests appeared to be much more dependent on sample complexity as compared to SpC analysis. We speculate that increasing complexity of MS1 spectra in our metaproteomes leads to mis-assignment of AUCs for specific mass peaks due to high peak density in the MS1 spectra.

Based on our evaluation, we recommend the analysis strategies for metaproteomics data in Table 3. Additionally, well-performing data pretreatment-test combinations are given in the upper parts of Figure 4 and in Figure 5. We provide the reproducible code, including the input files, and a readme file on how to execute the code with this study.

**Table 3:** Recommendation for metaproteomics statistical approaches based on our analyses. Please note that ranking (+++: high, ++: medium, +: low) is relative to each other.

|  | **Bayesian**<br>**regression**<br>**(brms)** | **edgeR** | **limma** | **Linear**<br>**model**<br>**(lm)** | **MaAsLin2** | **Random**<br>**forest**<br>**(rf)** |
| --- | --- | --- | --- | --- | --- | --- |
| Recommended<br>for/applicable to | SpC, AUC | SpC | SpC | SpC,<br>AUC | SpC, AUC | SpC, AUC |
| Data pretreatment | SpC:<br>gauss_chip09<br>AUC:<br>sn_chiPo9 | QL_RLE | voom | SpC:<br>logTSS<br>AUC:<br>sqrtTSS | SpC: astTSS<br>AUC: astTSS | SpC:<br>chiPo9_med<br>AUC: TMM |
| FDR control | +++ | +++ | +++ | +++ | +++ | ++ |
| Sensitivity (TPR) | +++ | ++ | ++ | + | + | ++ |
| Specificity (TNR) | + | ++ | ++ | ++ | +++ | +++ |
| Robust for large<br>community<br>changes/comparison<br>with low-abundant<br>community<br>members | ++ | + | ++ | + | ++ | ++ |
| Computational<br>demand | +++ | ++ | + | + | ++ | +++ |

One decisive factor in test choice might be the amount of computational resources needed or available to perform statistical testing. Computational speed varies widely, with limma (for SpC) and linear models being especially fast and feasible on desktop computers and laptops. EdgeR (for SpC) and MaAslin2 are suitable for desktop computers, but more intensive in terms of computational resources required. Using Bayesian regression, or random forests, requires more computational power and can usually only be executed feasibly on a high-performance desktop computer or server.

It is of note that all of the statistical inferences in our metaproteomics dataset find relative (and by no means absolute) differences between conditions and that the underlying metaproteomics quantification itself is relative, not absolute. This means that if we find a protein to be significantly more abundant in condition 1 as compared to condition 2, that same protein might still be more abundant on an absolute scale in condition 2, if overall more protein biomass is present in condition 2. It will be interesting to see whether and how recent approaches to estimate absolute microbial abundances from relative abundances via machine learning for large-scale datasets [44], or to model uncertainties/prior knowledge about the underlying total microbial community size explicitly [45, 46], can be adapted for metaproteomics and for environmental-omics studies.

In addition to the taxon-specific analyses of protein group abundances considered here, additional approaches are worth considering in a metaproteomics data analysis depending on the biological question addressed. For example, summing of protein group abundances for homologous proteins with the same functional annotation (isoforms), with and without considering taxonomic origin, can be useful for comparisons of overall (microbial) community function between different conditions. Along the same vein, protein abundances can also be summed for specific taxonomic groups or for complete metabolic pathways prior to statistical analysis.

In our study we focused on tests to compare two conditions. Such pairwise comparisons are also often at the core of more complex experimental setups. We did not evaluate approaches for longitudinal or paired/grouped designs (for the importance of taking grouped data into account, see, e.g., Vorland et al. 2025), but linear mixed models and approaches based on them, as well as random forests [48] can be adapted to accommodate grouped data.

Additionally, we want to stress that any statistic relies on an appropriate experimental design, and clearly formulated hypotheses [49]. Such a design needs to take into account, e.g., a sufficient number of replicates, appropriate controls, and mass spectrometry measurements in randomized blocks [50]. It is fundamental to think about how to analyze the data from the beginning, and to take into account whether to normalize at the organism level (see Methods and Kleiner 2017).

Our statistical performance evaluation framework is directly extendable to other metaproteomics data types such as DIA data, and other meta-omics approaches. DIA has been reported to give more reproducible protein identification results for metaproteomics [52], and might thus alleviate issues with data sparsity. At the same time, DIA, as well as feature mapping/match between runs [40, 53] can also introduce additional issues, e.g., potential mismatches in the case of match between runs [54], and database search issues [52], which still need to be addressed.

The evaluations of over 70 SpC and over 40 AUC statistical analysis methods, resulting recommendations, corresponding R code, and metaproteomics data we presented here will help microbiome scientists using metaproteomics to make informed choices about experimental design and statistics. The approaches we tested and recommend here are adaptable to different experimental designs. Using our results and our performance testing framework, researchers can move forward with analyzing their own meta(proteomics) data, using our provided code as a basis (see Availability of data and material). Moreover, researchers interested in developing and testing their own statistical approaches can use our publicly available data, and our evaluation code, for direct comparisons with the test strategies presented here.

## Supplementary Files

Supplementary Text incl. Supplementary Table 3 and Supplementary Figures 1-5 Supplementary Tables 1,2, 4-6

## Declarations

### Ethics approval and consent to participate

NC State’s Institutional Animal Care and Use Committee approved all experimental activity involving conventional and gnotobiotic mice (Protocol # 18-034-B and 18-165-B). No human data or tissue was used in this study.

### Consent for publication

Not applicable.

### Availability of data and material

Metagenomic sequencing data can be accessed via the ENA accession number PRJEB105044; fastq files can be directly accessed via http://ftp.sra.ebi.ac.uk/vol1/run/ERR159/ERR15997335/. All mass spectrometry proteomics data and results were deposited to the ProteomeXchange Consortium (http://proteomecentral.proteomexchange.org) via the PRIDE repository with the data set identifier PXD045390. Reviewers can currently access the repository using the following details: Username; Password: FwNk89lz.

We provide reproducible code, protein group quantification files, and an explanation of how to use the code via git (https://git.uni-greifswald.de/hinzket/Stats_Metaprot), as well as a fixed record via zenodo (DOI: 10.5281/zenodo.17880379).

## Author’s contributions

T.H. and M.K. conceived study and designed defined metaproteomes; T.H. performed all statistical analyses, generated figures, and wrote initial manuscript. B.J.K. generated defined metaproteome samples together with S.V., A.K., J.A.B.-R., performed metaproteomic sample preparation, measurement, and searches, generated figures. M.K. and B.J.K. gave input on statistical analyses. M.K. and P.W. mentored and supervised involved trainees. T.H., M.K., P.W., and B.J.K. acquired funding. All authors contributed to the manuscript.

## Acknowledgements

We thank Michael Greenacre for help with the chiPower transformation, and Philipp Adämmer for input on XGBoost. Elisa Kasbohm and Volkmar Liebscher provided valuable discussions on statistics. We thank Heather Maughan for valuable comments on the manuscript.

All LC-MS/MS measurements were made in the Molecular Education, Technology, and Research Innovation Center (METRIC) at North Carolina State University.

## References

1. Kleiner M. Metaproteomics: Much More than Measuring Gene Expression in Microbial Communities. mSystems. 2019;4:e00115–19. 10.1128/mSystems.00115-19.

2. Van Den Bossche T, Armengaud J, Benndorf D, Blakeley-Ruiz JA, Brauer M, Cheng K, et al. The microbiologist’s guide to metaproteomics. iMeta. 2025;4:e70031. 10.1002/imt2.70031.

3. Wilmes P, Bond PL. The application of two-dimensional polyacrylamide gel electrophoresis and downstream analyses to a mixed community of prokaryotic microorganisms. Environ Microbiol. 2004;6:911–20. 10.1111/j.1462-2920.2004.00687.x.

4. Blakeley-Ruiz JA, McClintock CS, Shrestha HK, Poudel S, Yang ZK, Giannone RJ, et al. Morphine and high-fat diet differentially alter the gut microbiota composition and metabolic function in lean versus obese mice. ISME Commun. 2022;2:66. 10.1038/s43705-022-00131-6.

5. Blakeley-Ruiz JA, Bartlett A, McMillan AS, Awan A, Walsh MV, Meyerhoffer AK, et al. Dietary protein source alters gut microbiota composition and function. ISME J. 2025;19:wraf048. 10.1093/ismejo/wraf048.

6. Levi Mortera S, Marzano V, Rapisarda F, Marangelo C, Pirona I, Vernocchi P, et al. Metaproteomics reveals diet-induced changes in gut microbiome function according to Crohn’s disease location. Microbiome. 2024;12:217. 10.1186/s40168-024-01927-5.

7. Mueller RS, Dill BD, Pan C, Belnap CP, Thomas BC, VerBerkmoes NC, et al. Proteome changes in the initial bacterial colonist during ecological succession in an acid mine drainage biofilm community. Environ Microbiol. 2011;13:2279–92. 10.1111/j.1462-2920.2011.02486.x.

8. Bergauer K, Fernandez-Guerra A, Garcia JAL, Sprenger RR, Stepanauskas R, Pachiadaki MG, et al. Organic matter processing by microbial communities throughout the Atlantic water column as revealed by metaproteomics. Proc Natl Acad Sci. 2018;115. 10.1073/pnas.1708779115.

9. Abbondio M, Tanca A, De Diego L, Sau R, Bibbò S, Pes GM, et al. Metaproteomic assessment of gut microbial and host functional perturbations in *Helicobacter pylori*-infected patients subjected to an antimicrobial protocol. Gut Microbes. 2023;15:2291170. 10.1080/19490976.2023.2291170.

10. Gruber-Vodicka HR, Leisch N, Kleiner M, Hinzke T, Liebeke M, McFall-Ngai M, et al. Two intracellular and cell type-specific bacterial symbionts in the placozoan *Trichoplax* H2. Nat Microbiol. 2019;4:1465–74. 10.1038/s41564-019-0475-9.

11. Langley SR, Mayr M. Comparative analysis of statistical methods used for detecting differential expression in label-free mass spectrometry proteomics. J Proteomics. 2015;129:83–92. 10.1016/j.jprot.2015.07.012.

12. Wolski WE, Nanni P, Grossmann J, d’Errico M, Schlapbach R, Panse C. prolfqua: A Comprehensive R-Package for Proteomics Differential Expression Analysis. J Proteome Res. 2023;22:1092–104. 10.1021/acs.jproteome.2c00441.

13. Yang Y, Cheng J, Wang S, Yang H. StatsPro: Systematic integration and evaluation of statistical approaches for detecting differential expression in label-free quantitative proteomics. J Proteomics. 2022;250:104386. 10.1016/j.jprot.2021.104386.

14. Zhu Y, Orre LM, Zhou Tran Y, Mermelekas G, Johansson HJ, Malyutina A, et al. DEqMS: A Method for Accurate Variance Estimation in Differential Protein Expression Analysis. Mol Cell Proteomics. 2020;19:1047–57. 10.1074/mcp.TIR119.001646.

15. Malinowska A, Kistowski M, Bakun M, Rubel T, Tkaczyk M, Mierzejewska J, et al. Diffprot — software for non-parametric statistical analysis of differential proteomics data. J Proteomics. 2012;75:4062–73. 10.1016/j.jprot.2012.05.030.

16. Pursiheimo A, Vehmas AP, Afzal S, Suomi T, Chand T, Strauss L, et al. Optimization of Statistical Methods Impact on Quantitative Proteomics Data. J Proteome Res. 2015;14:4118–26. 10.1021/acs.jproteome.5b00183.

17. Ramus C, Hovasse A, Marcellin M, Hesse A-M, Mouton-Barbosa E, Bouyssié D, et al. Benchmarking quantitative label-free LC–MS data processing workflows using a complex spiked proteomic standard dataset. J Proteomics. 2016;132:51–62. 10.1016/j.jprot.2015.11.011.

18. Li M, Gray W, Zhang H, Chung CH, Billheimer D, Yarbrough WG, et al. Comparative Shotgun Proteomics Using Spectral Count Data and Quasi-Likelihood Modeling. J Proteome Res. 2010;9:4295–305. 10.1021/pr100527g.

19. Calgaro M, Romualdi C, Waldron L, Risso D, Vitulo N. Assessment of statistical methods from single cell, bulk RNA-seq, and metagenomics applied to microbiome data. Genome Biol. 2020;21:191. 10.1186/s13059-020-02104-1.

20. Jonsson V, Österlund T, Nerman O, Kristiansson E. Statistical evaluation of methods for identification of differentially abundant genes in comparative metagenomics. BMC Genomics. 2016;17:78. 10.1186/s12864-016-2386-y.

21. Nearing JT, Douglas GM, Hayes MG, MacDonald J, Desai DK, Allward N, et al. Microbiome differential abundance methods produce different results across 38 datasets. Nat Commun. 2022;13:342. 10.1038/s41467-022-28034-z.

22. Weiss S, Xu ZZ, Peddada S, Amir A, Bittinger K, Gonzalez A, et al. Normalization and microbial differential abundance strategies depend upon data characteristics. Microbiome. 2017;5:27. 10.1186/s40168-017-0237-y.

23. Chen Y, Chen L, Lun ATL, Baldoni PL, Smyth GK. edgeR v4: powerful differential analysis of sequencing data with expanded functionality and improved support for small counts and larger datasets. Nucleic Acids Res. 2025;53:gkaf018. 10.1093/nar/gkaf018.

24. Love MI, Huber W, Anders S. Moderated estimation of fold change and dispersion for RNA-seq data with DESeq2. Genome Biol. 2014;15:550. 10.1186/s13059-014-0550-8.

25. Ritchie ME, Phipson B, Wu D, Hu Y, Law CW, Shi W, et al. limma powers differential expression analyses for RNA-sequencing and microarray studies. Nucleic Acids Res. 2015;43:e47–e47. 10.1093/nar/gkv007.

26. Breiman L. Random Forests. Mach Learn. 2001;45:5–32.

27. Gloor GB, Macklaim JM, Pawlowsky-Glahn V, Egozcue JJ. Microbiome Datasets Are Compositional: And This Is Not Optional. Front Microbiol. 2017;8:2224. 10.3389/fmicb.2017.02224.

28. Lazar C, Gatto L, Ferro M, Bruley C, Burger T. Accounting for the Multiple Natures of Missing Values in Label-Free Quantitative Proteomics Data Sets to Compare Imputation Strategies. J Proteome Res. 2016;15:1116–25. 10.1021/acs.jproteome.5b00981.

29. Plancade S, Berland M, Blein-Nicolas M, Langella O, Bassignani A, Juste C. A combined test for feature selection on sparse metaproteomics data—an alternative to missing value imputation. PeerJ. 2022;10:e13525. 10.7717/peerj.13525.

30. Välikangas T, Suomi T, Elo LL. A systematic evaluation of normalization methods in quantitative label-free proteomics. Brief Bioinform. 2016;:bbw095. 10.1093/bib/bbw095.

31. Díaz-Uriarte R, Alvarez De Andrés S. Gene selection and classification of microarray data using random forest. BMC Bioinformatics. 2006;7:3. 10.1186/1471-2105-7-3.

32. Blakeley-Ruiz JA, Kleiner M. Considerations for constructing a protein sequence database for metaproteomics. Comput Struct Biotechnol J. 2022;20:937–52. 10.1016/j.csbj.2022.01.018.

33. R Core Team. R: A Language and Environment for Statistical Computing. R Found Stat Comput Vienna Austria. 2023.

34. Benjamini Y, Hochberg Y. On the Adaptive Control of the False Discovery Rate in Multiple Testing with Independent Statistics. J Educ Behav Stat. 2000;25:60–83.

35. Benjamini Y, Yekutieli D. The control of the false discovery rate in multiple testing under dependency. Ann Stat. 2001;29:1165–88.

36. Brodersen KH, Ong CS, Stephan KE, Buhmann JM. The balanced accuracy and its posterior distribution. 2010 Int Conf Pattern Recognit. 2010;:3121–4.

37. Sasaki Y. The truth of the F-measure. 2007.

38. Sitarz M. Extending F1 metric, probabilistic approach. 2022. 10.48550/arXiv.2210.11997.

39. Li Y, Ge X, Peng F, Li W, Li JJ. Exaggerated false positives by popular differential expression methods when analyzing human population samples. Genome Biol. 2022;23:79. 10.1186/s13059-022-02648-4.

40. Tyanova S, Temu T, Cox J. The MaxQuant computational platform for mass spectrometry-based shotgun proteomics. Nat Protoc. 2016;11:2301–19. 10.1038/nprot.2016.136.

41. Yerke A, Fry Brumit D, Fodor AA. Proportion-based normalizations outperform compositional data transformations in machine learning applications. Microbiome. 2024;12:45. 10.1186/s40168-023-01747-z.

42. Webb-Robertson B-JM, Wiberg HK, Matzke MM, Brown JN, Wang J, McDermott JE, et al. Review, Evaluation, and Discussion of the Challenges of Missing Value Imputation for Mass Spectrometry-Based Label-Free Global Proteomics. J Proteome Res. 2015;14:1993–2001. 10.1021/pr501138h.

43. Schurch NJ, Schofield P, Gierliński M, Cole C, Sherstnev A, Singh V, et al. How many biological replicates are needed in an RNA-seq experiment and which differential expression tool should you use? RNA. 2016;22:839–51. 10.1261/rna.053959.115.

44. Nishijima S, Stankevic E, Aasmets O, Schmidt TSB, Nagata N, Keller MI, et al. Fecal microbial load is a major determinant of gut microbiome variation and a confounder for disease associations. Cell. 2025;188:222–236.e15. 10.1016/j.cell.2024.10.022.

45. Gloor GB, Nixon MP, Silverman JD. Explicit Scale Simulation for analysis of RNA-sequencing count data with ALDEx2. NAR Genomics Bioinforma. 2025;7:lqaf108. 10.1093/nargab/lqaf108.

46. Nixon MP, Gloor GB, Silverman JD. Incorporating scale uncertainty in microbiome and gene expression analysis as an extension of normalization. Genome Biol. 2025;26:139. 10.1186/s13059-025-03609-3.

47. Vorland CJ, Golzarri-Arroyo L, Allison DB. A brief guide to statistical analysis of grouped data in preclinical research. Nat Metab. 2025;7:1301–4. 10.1038/s42255-025-01323-9.

48. Capitaine L, Genuer R, Thiébaut R. Random forests for high-dimensional longitudinal data. Stat Methods Med Res. 2021;30:166–84. 10.1177/0962280220946080.

49. Wagner MR, Kleiner M. How thoughtful experimental design can empower biologists in the omics era. Nat Commun. 2025;16:7263. 10.1038/s41467-025-62616-x.

50. Oberg AL, Vitek O. Statistical Design of Quantitative Mass Spectrometry-Based Proteomic Experiments. J Proteome Res. 2009;8:2144–56. 10.1021/pr8010099.

51. Kleiner M. Normalization of metatranscriptomic and metaproteomic data for differential gene expression analyses: The importance of accounting for organism abundance. 2017. 10.7287/peerj.preprints.2846v1.

52. Rajczewski AT, Blakeley-Ruiz JA, Meyer A, Vintila S, McIlvin MR, Van Den Bossche T, et al. Data-Independent Acquisition Mass Spectrometry as a Tool for Metaproteomics: Interlaboratory Comparison Using a Model Microbiome. PROTEOMICS. 2025;25:e202400187. 10.1002/pmic.202400187.

53. Yu F, Haynes SE, Nesvizhskii AI. IonQuant Enables Accurate and Sensitive Label-Free Quantification With FDR-Controlled Match-Between-Runs. Mol Cell Proteomics. 2021;20:100077. 10.1016/j.mcpro.2021.100077.

54. Lim MY, Paulo JA, Gygi SP. Evaluating False Transfer Rates from the Match-between-Runs Algorithm with a Two-Proteome Model. J Proteome Res. 2019;18:4020–6. 10.1021/acs.jproteome.9b00492.

