## Supplementary Text for "Comprehensive evaluation of statistical approaches for differential metaproteomics"

### Supplementary Material and Methods

#### Cultivation of microorganisms

*Escherichia coli* strain HS was grown on either LB complex medium [1] or M9 minimal medium [2] with 1% glucose at 37 °C until the bacteria reached exponential phase; 3 h and 20 min for LB and 4 h and 30 min for M9. Some 1 L of cultures were grown in quadruplicates for each condition. Samples were pelleted by centrifugation at 8,000 g for 30 min. Supernatant was removed and pellets were resuspended in 100 mL of PBS. Suspended pellets were aliquoted into 64 tubes in 1 mL aliquots. Aliquots were centrifuged at 17,000 x g for 10-15 min. Supernatant was removed and frozen at -80 °C.

*Bacterioides thetaiotaomicron* was grown in degassed serum bottles in meat broth [3] at 37 °C in quadruplicates overnight until turbid. 1.5 ml aliquots were removed and centrifuged at 13,000 x g for 5 min. Pellets were resuspended in 1 ml sterile PBS buffer and pelleted again at 13,000 x g for 5 min. Pellets were immediately stored at -80 °C.

*Thermus thermophilus* strain HB27 was grown in quadruplicates at either 70 °C (high temperature) or 55 °C (low temperature). Cultures were grown in 1 L TSB (Difco) supplemented with 0.4% yeast extract and 0.3% NaCl, for 24-48 h and harvested at 15,000 x g for 20 min at 4 °C. Pellets were resuspended in 100 ml sterile PBS buffer, aliquoted into 1.7 ml Eppendorf tubes and centrifuged again at 15,000 x g for 5 min at 4 °C. Pellets were immediately stored at -80 °C.

*Chlamydomonas reinhardii* CC-3403 was grown in TAP-medium [4] supplemented with 267 mM arginine. Cultures were grown at room temperature under either 270 µE (high light) or 27 µE (low light) with a 12:12 light:dark cycle. Some 1 L of cultures were grown for 72-96 h in quadruplicates for each condition and harvested at 10,000 x g for 20 min at 4 °C. Pellets were resuspended in 110 ml sterile PBS, aliquoted into 1.7 ml Eppendorf tubes and centrifuged again at 10,000 x g for 5 min at 4 °C. Pellets were immediately stored at -80 °C.

*Rhizobium leguminosarum* strains RL3841 and VF39 were grown in 1L YEM-broth [5]. Each strain was grown in quadruplicates at 28 °C for 48 h. Cultures were harvested at 15,000 x g for 20 min at 4 °C. Pellets were resuspended in 110 ml sterile PBS, aliquoted into 1.7 ml Eppendorf tubes and centrifuged again at 15,000 x g for 5 min at 4 °C. Pellets were immediately stored at -80 °C.

#### Metaproteomics database generation

Databases for the different mixes were built from combining building blocks as follows: For mixes M01 to M06 based on the matrix made of standard mouse faecal pellets, the protein sequence database was constructed by combining the clustered microbial translated metagenome (see below section "Mouse metagenome sequencing and analysis"), the clustered mouse proteome, the clustered mouse diet proteome, and contaminants. The resulting matrix database was clustered at 99% using CD-HIT-2D [6] with the database of clustered microbial isolate genomes to remove sequences from the matrix database that were very similar to sequences from the pure cultures. The resulting matrix-specific protein sequences were concatenated with all microbial isolate protein sequences to generate the final database for the mixes M01 to M06.

For the mixes M07 to M11 based on the sterile corn root matrix, the matrix proteome database consisted of the clustered corn proteome and the contaminants. Like the mouse proteome, the resulting matrix database was clustered at 99% using CD-HIT-2D with the clustered microbial isolate database to remove sequences from the matrix database that were very similar to sequences from the pure culture isolates. The resulting matrix-specific protein sequences were concatenated with the microbial isolate proteomes to generate the final database for the mixes M07 to M11.

For the mixes M12 and M13 based on the gnotobiotic mouse matrix, the matrix protein sequence database consisted of the clustered proteins of the reduced microbial community, the clustered mouse proteome (see above), the clustered diet proteins and the contaminants (see above). The resulting matrix database was clustered at 99% using CD-HIT-2D with the clustered isolates to remove sequences from the matrix database that were very similar to sequences from the pure culture isolates. The resulting matrix-specific sequences were concatenated with the pure culture proteomes to generate the final database for the mixes M12 to M13.

#### Defined metaproteome protein sample generation

##### Pure culture peptide generation

Pellets of pure cultures were used for protein extraction and digested into peptides using the filter-aided sample preparation (FASP; Wiśniewski *et al.*, 2009; Blakeley-Ruiz *et al.*, 2025) with few modifications as follows. Briefly, for each pellet, ~300 µL of SDT lysis buffer (4% (w/v) SDS, 100 mM Tris-HCl pH 7.6, 0.1 M DTT) was added. Some samples underwent bead beating using lysing matrix E tubes (MP Biomedicals) for 5 cycles of 45 s at 6,45 m/s speed and 1 min pause between cycles (see Supplementary Table 1 for details). All samples were heated to 95 °C for 10 min and centrifuged at 21,000 x g for 5 min prior being transferred to several FASP filters (60 µL/filter). We mixed 60 µl of the lysate with 400 µL of UA solution (8 M urea in 0.1 M Tris/HCl pH 8.5) onto a modified PES 10 kDa 500 µL filter unit (VWR International) and centrifuged at 14,000 x g for 30 min. The filters were washed using 200 µL of UA solution and centrifuged at 14,000 g for 40 min followed by incubation with 100 µL IAA (0.05 M iodoacetamide in UA solution) for 20 min and centrifugation at 14,000 g for 20 min. Filters were then washed three times with 100 µL of UA buffer and 3 times using 100 µL of ABC (50 mM Ammonium Bicarbonate) buffer. The proteins were digested into peptides by adding 0.8 µg of MS grade trypsin (Thermo Scientific Pierce) solubilized in 40 µL of ABC buffer to the filters and incubating for 16 h in a wet chamber at 37°C. Peptides were eluted by centrifugation at 14,000 x g for 20 min followed by another elution using 50 µL of 0.5 M NaCl and centrifugation at 14,000 x g for 20 min. Extracted peptides from the same samples extracted on different filters were combined into ~800-1000 µL of resulting peptides, and 0.4 to 0.5 ml of 0.1% formic acid (FA) was added prior to desalting using C18 Sep-Pak Light cartridges (Waters) and syringes. Briefly, the Sep-Pak cartridge was washed and primed with 5 ml of wash solution (70% ACN, 0.1% FA) followed by 5 ml of 0.1% FA solution.

The sample was loaded in three steps on the cartridge, rinsed with 5 ml of wash solution and eluted slowly into two tubes using 1.5 ml of elution solution (90% ACN, 0.1% FA). The samples were concentrated and depleted of acetonitrile with a first vacuum centrifugation step to ~300 µL and solvent was exchanged by adding 600 µL of 0.1% FA solution followed by a second vacuum centrifugation step until a final volume of ~300µL was reached. Resulting peptide concentrations were measured in triplicate using the Pierce Micro BCA assay (Thermo Scientific Pierce) according to the manufacturer's instructions.

##### Matrix peptide generation

For the conventional mouse stool matrix, fecal pellets from a previous study [7] were gathered from four different cages and homogenized together to form a representative sample of ~1.6 g fecal pellet, from which 500 mg were used for metagenomics analysis. Proteins and peptides were extracted and processed similarly to the pure cultures using the FASP protocol (see above). For obtaining corn roots, *Zea mays* seeds were sterilized using the protocol described in Parnell *et al.* (2024) by immersing the seeds in 70% ethanol for 3 min followed by immersion in 2% sodium hypochlorite for 3 min. Seeds were rinsed in de-ionized $H_2O$ ten times before being placed individually in sterile WhirlPak bags (Nasco) containing 90 ml of sterile clay (Pro's Choice Rapid Dry, OIL-DRI) and watered with 90 ml sterile 0.5x Murashige-Skoog media. Bags were sealed with Aeraseal (Millipore Sigma) and plants were grown for 13 days at 25 °C with 16:8 hours light:dark cycle. Roots were harvested, chopped and frozen at -80 °C until further processing. For the corn roots and gnotobiotic mice samples [10], the FASP protocol was not providing sufficient yield and the extractions were performed using the S-Trap mini MS sample prep kit (Protifi) instead. The original S-Trap manufacturer's protocol uses different reductant and alkylator as compared to the FASP protocol, which would have precluded mixing of the samples. Therefore, the S-Trap protocol was modified to use DTT and IAA and was applied as follows. For both matrices, 100 mg of material was mixed with 1 mL of SDS lysis buffer (5% (w/v) SDS, 100 mM Tris-HCl pH 7.6, 0.1 M DTT), bead-beated using lysing matrix E tubes (MP Biomedicals; 5 cycles of 45 s at 6,45 m/s speed and 1 min pause between cycles), heated to 95 °C for 10 min, and centrifuged at 21,000 x g for 5 min. Samples were then aliquoted into 140 µL of sample. Reduction was performed by adding 6.4 µL of 500 mM DTT and incubating at 95 °C for 10 min. Subsequently, samples were alkylated by adding 12.7 µL of 500 mM IAA and incubating in the dark at room temperature for 30 min. Samples were then acidified using 16 µL of 12% phosphoric acid solution (to reach pH<1). Finally, 1050 µL of binding/wash buffer (100 mM TEAB in 90% methanol) was mixed with the samples to form protein colloids, loaded (two times 600 µL) onto S-Trap columns and centrifuged at 4,000 x g for 30 s. The column was washed three times with 400 µL of binding/wash buffer and centrifuged at 4,000 x g for 30 s, followed by a final centrifugation at 4,000 x g for 1 min to fully remove the buffer. Proteins were then digested into peptides by adding 0.8 µg of MS grade trypsin (Thermo Scientific Pierce) solubilized in 40 µL of digestion/elution buffer (50 mM TEAB in water) to the filters and incubating for 16 h in a wet chamber at 37 °C. Peptides were eluted by adding 80 µL of digestion/elution buffer, incubation 10 min at 37 °C and centrifugation at 4,000 x g for 1 min; followed by two more elutions by adding 80 µL of elution buffer 2 (0.2% FA) and centrifugation at at 4,000 x g for 1 min, and finally adding 80 µL of elution buffer 3 (50% ACN, 0.2% FA) and centrifugation at at 4,000 x g for 1 min. The resulting 240 µL were concentrated and depleted of acetonitrile using vacuum centrifugation until a volume of ~60 µL was reached. Desalting of the peptides and their concentration measurement were performed as for the pure cultures (see above).

##### Defined metaproteome production

Following peptide quantification, the matrices and all quadruplicates of the pure cultures were diluted to 250 ng/µL and the dilutions were measured again in triplicate using the Pierce Micro BCA assay (Thermo Scientific Pierce) to ensure that the correct concentration was achieved. Aliquots were generated at all steps to subject samples to freeze-thaw cycles only once per preparation step. Peptides extracted from the different isolates and conditions were then mixed in different proportions with the respective matrix peptides to produce the various defined metaproteomes (see Main Text, Figure 1 and Supplementary Table 4).

#### Mouse metagenome sequencing and analysis

To generate generic mouse stool matrix protein data for the metaproteomics database, two times 250 mg of fecal pellets were used for DNA extraction with the DNeasy PowerSoil Pro Kit following manufacturer's instructions except that bead beating was used instead of vortexing (3.1 m/s for 3 cycles of 30 sec. with 1 min of cooling on ice in between each cycle) in 2 ml bead beating tubes (Lysing Matrix E, MP Biomedicals) using a Bead Ruptor Elite 24 (Omni International). The extracted DNA was eluted with 60 µL of solution C6, merged into one tube and DNA concentration of the combined extractions was assessed using a DS-11 FX+ Spectrophotometer (Denovix) with the Qubit™ dsDNA High Sensitivity Assay Kit (Invitrogen). Metagenomic DNA was submitted to the North Carolina State Genomic Sciences Laboratory (Raleigh, NC, USA) for Illumina library construction and sequencing to produce ~400 M 150 bp paired-end reads. Library construction was performed using an Illumina TruSeq Nano Library kit according to manufacturer's instructions. Libraries were sequenced on an Illumina NextSeq 500 sequencer.

Removal of the Illumina Truseq3 adapters and quality trimming of the raw reads was performed using Trimmomatic v0.39 [11], removing all bases on the 3′-end with a Phred score lower than 20 (if present) and excluding all reads shorter than 40 bp. The trimmed reads were assembled using MEGAHIT v1.2.9 [12] with default parameters, minimum and maximum k-mer sizes of 25 and 99 respectively and a k-mer increment of 4. The trimmed reads were mapped against the resulting assembly contigs using the Burrows-Wheeler Aligner with maximal exact matches (BWA-MEM; Li and Durbin, 2009) algorithm requiring 100% identity. The coverage was used to bin the assembly contigs with several binners, namely, MetaBAT v2.2.15 [14], Maxbin 2.2.7 [15], SemiBin v1.0.3 [16], Binny v2 [17] and MetaDecoder v1.0.13 [18] with default parameters. The bins from the different tools were then refined with DAS Tool v1.1.4 [19] using a score threshold of 0.7 and keeping both the contigs belonging to refined bins and the unbinned fraction. Open reading frames (ORFs) were then predicted on both fractions using Prodigal v2.6.3 [20] and the resulting ORFs were used for database generation.

#### Protein quantification data preprocessing

##### Data filtering and imputation

We analyzed differential protein abundance in samples by comparing one defined metaproteome vs. another, within the same matrix. These defined metaproteomes simulate independent experimental conditions, environmental conditions, spatial sampling points, or other groups of samples to be compared. We used master proteins for our analysis. Master proteins are representative sequences chosen for protein groups, where protein groups are one or multiple protein sequences which share a set of peptide-spectrum matches [21, 22]. Selection of master proteins for specific protein groups can differ in pure cultures vs. defined metaproteomes, leading to empty matches and thereby inaccurate comparisons. We addressed this issue by matching on a per-protein level: only one protein of the pure culture protein group needed to be present in a defined metaproteome protein group for matching. Sparsity in the dataset was reduced by filtering the SpC data for most implementations to retain only proteins that had at least 5 PSMs in at least 2 out of 8 samples. AUC data were initially filtered to arrive at approximately the same number of proteins as for SpC, to make the two quantification approaches comparable. The effect of filtering was subsequently assessed with selected statistical methods (see Supplementary Results and Supplementary Figure 1). We imputed missing values with 0 or, where necessary, with a small constant (1/5th of the smallest value in the dataset). For random forests that use clr transformed data, we also tested the zcompositions count imputation [23].

##### Normalization and transformation

For normalization, which is done to correct for differences in overall quantification (sum of AUC or PSMs) between mass spectrometry runs, as well as unequal abundances of the species to be compared between samples in the case of species-level normalization [24], we used total sum scaling (TSS), cumulative sum scaling (CSS), and (organism-level) normalized spectral abundance factors ((org)NSAF; [25, 26]).

Additionally, we used the trimmed mean of M-values (TMM) normalization [27], and its new variant TMM with singleton pairing (TMMwsp; [28]), which was developed for more robust analysis of low-abundant variables, as well as the relative log expression (RLE; Anders and Huber, 2010). The voom normalization for limma [30] uses the mean-variance trend for precision weight estimation. These weights are then used in subsequent analysis. DeqMS [31], another limma normalization method, which was explicitly developed for proteomics data, models the prior variance per group depending on the number of PSMs or peptides. Additionally, we employed variance stabilization normalization (vsn; [32], which aims to make the standard deviation independent of the mean.

We used different transformations to make the data, or the residuals of a linear modelling approach, normally distributed. We used log2, logit, sqrt and arcsine square root (ast) transformation. The sqrt of TSS is also called Hellinger transformation [33]. Additionally, we used the centered log ratio (clr) transformation, which was proposed to remedy the issue of compositionality [34]. More recently, the chiPower (chiP) transformation was proposed as an alternative to clr, which does not require zero replacement [35]. For chiP transformation, protein group abundances are first raised to power lambda, then devided by the sample total, chi-square standardized (divided by the square root of the mean of that protein across samples), BoxCox-transformed and centered. We tested different values of lambda. See Table 3 of the Main Text for combinations of normalizations/transformations and tests.

#### Statistical analyses

##### Regression-based tests

*(Bayesian) (Generalized) linear mixed model*

For (generalized) linear regression, we used a random effects only-model:

value ~ (1 + condition | protein)

We implemented Gaussian regression with transformed data, as well as Poisson and negative binomial regression in lme4 v. 1.1.34 [36]. Significant differences were defined as differences where the interval of the slope of the random effect ± 1.96 the standard deviation did not include 0 (i.e., where the 95% confidence interval of the random slope was strictly positive or strictly negative). Few linear model evaluations produced boundary fits.

For the Bayesian approach, we used brms [37] with the same model formula as above. We used 3 chains, 6000 iterations with 2000 of these assigned to warmup, and chain thinning of 10. We assigned significance if the 2.5 to 97.5% credibility interval of the conditional random effect excluded zero. We tested Gaussian, as well as exponentially modified Gaussian, students, and skew-normal distributions, with different normalizations. We mostly used the specified default priors, aside from a model with Gaussian distribution and chiP (lambda=0.9) normalization and transformation, where we also adapted priors. In this case, we modelled, with a Student's t-distribution, the intercept (3,0,0.1), residual standard deviation sigma (3,0,1), and standard deviation of the group-level (random) effect (3,0,2). For SpC data, brms evaluations in matrix 3 produced low effective sample sizes (ESS), with the exception of brms_gauss_chiP09_prior, i.e., adaptation of priors was necessary to achieve sufficiently fast convergence.

*Corncob*

The R package corncob [38] uses a beta-binomial regression model, which was developed for differential abundance analysis of taxa in microbial communities. It explicitly takes overdispersion into account, thus allowing for testing differential relative abundance together with differential variance. For that, it uses a logit link, and directly uses absolute abundances and total counts in the model, instead of normalizing beforehand. FDR correction is implemented in the algorithm. We used corncob v. 0.3.1 [39].

*DESeq2 (negative binomial)*

DESeq2 [40], originally developed for RNA-Seq data, models data with a negative binomial distribution, which can be formulated as a mixture of gamma and Poisson distributions. It normalizes for different sequencing depths and uses a logarithmic link in its generalized linear model. A shrinkage of dispersion estimates within groups is also employed, with stronger shrinkage for lower information-containing variables. DESeq2 has FDR calculation implemented.

*edgeR 4.0*

EdgeR [28] uses a negative binomial distribution for counts, and can accommodate different experimental designs via generalized linear models. It employs empirical Bayes moderation for outlier treatments, i.e., the dispersions are estimated from the data, and variations moderated towards the common trend [28]. The newest version edgeR v4.0 uses unbiased quasi-likelihood dispersion estimates (QLE) also for small counts. We used the FDR calculation as implemented.

*Limma*

Limma [41], originally developed for microarray data (= continuous intensity data) and extended to accommodate RNAseq data (= count data), uses a variable-wise linear model with global parameters to share information across genes and samples. An empirical Bayes moderation is used to moderate the residual variance. Limma has FDR calculation implemented.

*MaAsLin2*

MaAsLin2 [42] uses generalized linear mixed models, with the option to use different model specifications, as well as normalizations and transformations. It accommodates different experimental designs and the inclusion of metadata. We used the implemented FDR value calculation.

#### Ensemble machine learning

*Random forest*

For random forests [43], we used the variable importance measure of Janitza et al. [44]. Additionally, we aggregated results over forests per comparison to increase stability. We used 500 forests and 75,000 trees per forest when not normalizing at the species level, and 500 forests with 50,000 trees per forest when normalizing at the species level in the defined metaproteomes, as well as in the pure cultures. We assigned protein groups as significantly differentially abundant when the mode of the p-value density distribution of forests was below 0.05, or when the median of p-values of all forests for one comparison was below 0.05. Monotonic transformations like a log transfomation will not impact the random forest results, as the splits are the same. Some random forest p-value estimations were inaccurate due to too few non-negative values (the alternative Altmann approach for p-value estimation was prohibitively slow and therefore not used). Each random forest result was aggregated over forests as well as over trees to ameliorate p-value issues.

#### Null hypothesis testing

*Welch's t-test*

While the Student's t-test assumes normality and homogeneity of variances, Welch's t-test approximation does not require homogeneity of variances [45]. As Welch's t-test performs better type I error control with heterogenous variances and looses little power (if any) as compared to the Student's t-test, it is recommended to be used as default instead of the Student's t-test [46].

We calculated Benjamini-Hochberg corrected FDR [47, 48] values from p-values using adjust_pvalue in rstatix [49].

*Wilcoxon rank-sum test*

The Wilcoxon rank-sum test is based on ranking observations of both groups by their abundance and then summing these ranks per group. The Wilcoxon rank-sum test does not make assumptions about the data distribution and is thus a non-parametric test to compare two conditions [50]. FDR values were calculated from p-values using adjust_pvalue in rstatix [49].

##### Approaches not included

Some approaches were initially tested, but not included in the final analyses. XGBoost [51] needed prohibitively complex hyperparameter tuning in order to give meaningful results. ALDEx2 [52] is known to have low power [53], an observation we also made during testing. An explicit scale model, as very recently published [54], was not tested, and is not expected to ameliorate low power issues in our few-replicate setting, especially as it was noted that scale uncertainty inclusion decreases sensitivity [55]. Likewise, the experimental layout we describe here, i.e., few replicates per condition, does not lend itself to neural network-based machine learning approaches.

#### Software implementation

For versions of R 4.3.1 [56] packages used, please see Supplementary Table 3.

*Supplementary Table 3: R Packages and versions used in this study.*

| Package | Version | Reference |
|---|---|---|
| **General data reformatting** | | |
| dplyr | 1.1.4 | [57] |
| plyr | 1.8.9 | [58] |
| tidyr | 1.3.1 | [59] |
| tidyverse | 2.0.0 | [60] |
| **Statistics and helper functions for statistics** | | |
| brms | 2.22.0 | [37] |
| cmdstanr | 0.8.1 | [61] |
| compositions | 2.0-6 | [62] |
| corncob | 0.3.1 | [39] |
| DEqMS | 1.2.0 | [31] |
| DESeq2 | 1.40.2 | [40] |
| edgeR | 4.0.16 | [28] |
| irr | 0.84.1 | [63] |
| limma | 3.58.1 | [41] |
| lme4 | 1.1-36 | [36] |
| lmerTest | 3.1-3 | [64] |
| Maaslin2 | 1.16.0 | [42] |
| matrixTests | 0.2.3 | [65] |
| ranger | 0.16.0 | [66] |
| rstatix | 0.7.2 | [49] |
| TAF | 4.2.0 | [67] |
| tidymodels | 1.2.0 | [68] |
| zCompositions | 1.5.0-4 | [23] |
| **Parallel job execution** | | |
| doParallel | 1.0.17 | [69] |
| foreach | 1.5.2 | [70] |
| **Figure generation** | | |
| circlize | 0.4.16 | [71] |
| ComplexHeatmap | 2.18.0 | [72] |
| egg | 0.4.5 | [73] |
| gghalves | 0.1.4 | [74] |
| ggplot2 | 3.5.1 | [75] |
| patchwork | 1.1.3 | [76] |
| RColorBrewer | 1.1-3 | [77] |
| scico | 1.5.0 | [78]; color palette: batlow [79] |
| svglite | 2.1.1 | [80] |

### Supplementary Results and Discussion

#### Filtering increases true positives and impacts true negatives

We tested the effect of filtering by evaluating within-condition abundance changes, i.e., evaluations which are independent of comparing with the pure-culture proteomes, as a pure culture proteome ground truth would also be dependent on the filtering (and therefore would need to be evaluated for each filtering level). For SpC, we varied filter criteria between at least 1 PSM in at least 25% of samples (lowest stringency) up to at least 20 PSM in at least 50% of samples (highest stringency). For AUC, we varied between an intensity of at least 5.000.000 in at least 25% of samples (lowest stringency) to at least 200.000.000 in at least 50% of samples (highest stringency). Increasingly stringent filtering by the number of samples which need to contain a certain protein group abundance value and the protein group abundance value itself generally increased the percentage of true positives, but did not impact or decreased the percentage of true negatives (Supplementary Figure 1). Obviously, also the absolute number of protein groups in the comparison decreased with more stringent filtering. For random forests using SpC data, too-harsh filtering criteria caused the p-value calculation to fail, as no negative importances were found. In the case of AUC data, too-lenient filtering criteria led to brms and the random forest being prohibitively slow.

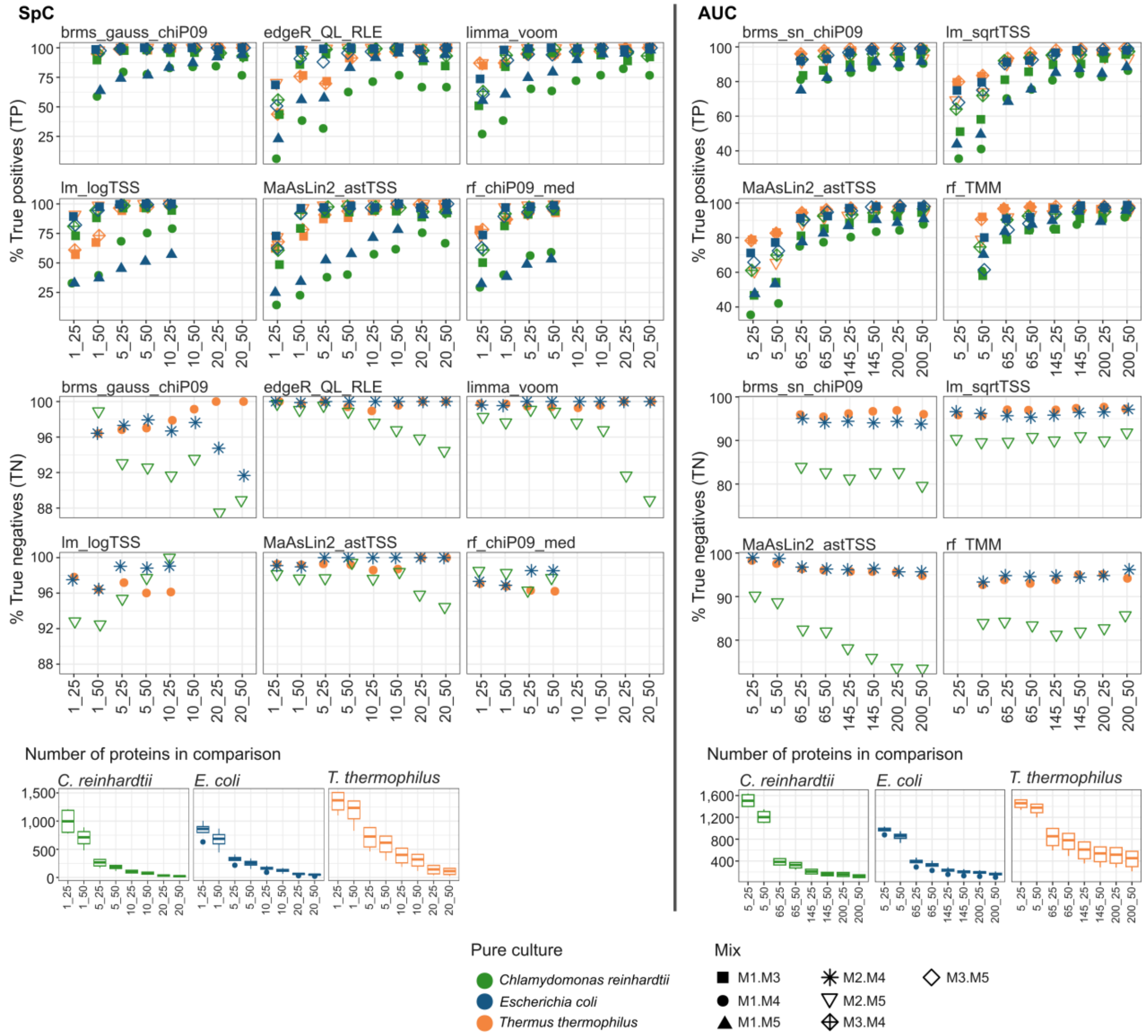


*Supplementary Figure 1: Effect of filtering on the percentage ot true positives and true negatives found for the respective comparison. For spectral count (SpC) data, numbers on the x axis indicate the minimum number of spectra required in a certain share of samples, e.g., 1_25 means that at least 1 PSM is required in at least 25% of the samples. For peptide intensity data (AUC), numbers on the x axis indicate the minimum intensity (times $10^6$) required in a certain share of samples, e.g., 5_25 means an intensity of 5.000.000 required in at least 25% of samples. Missing values indicate that the calculation was not possible, either because too many values were missing (rf for SpC data) or because the model evaluation got prohibitively slow due to a too-large dataset (brms and rf for AUC).*

#### Matrix complexity and abundance changes impact AUC data evaluation performance more than SpC data evaluation performance

Marked differences existed in P4 scores depending on abundances of organisms compared, but also depending on the organisms themselves, with *C. reinhardtii* producing consistently lower results (Supplementary Figures 1 and 5, Supplementary Tables 5e, 5h, 5i, 6e, 6h, 6i). While AUC comparisons performed better for the higher-abundant *T. thermophilus* and *E. coli* comparisons, their performance loss was relatively and absolutely higher than that of SpC-based comparisons for the other tests (see Main Text). The differential performance of SpC vs. AUC-based statistics in general might partially be caused by a match-between-runs transfer of peptide identifications and AUC quantifications, while SpC-based data does not use match-between-runs, concomitant with less consistent protein identifications especially for *C. reinhardtii* (Supplementary Figures 3, 4, and Main Text).

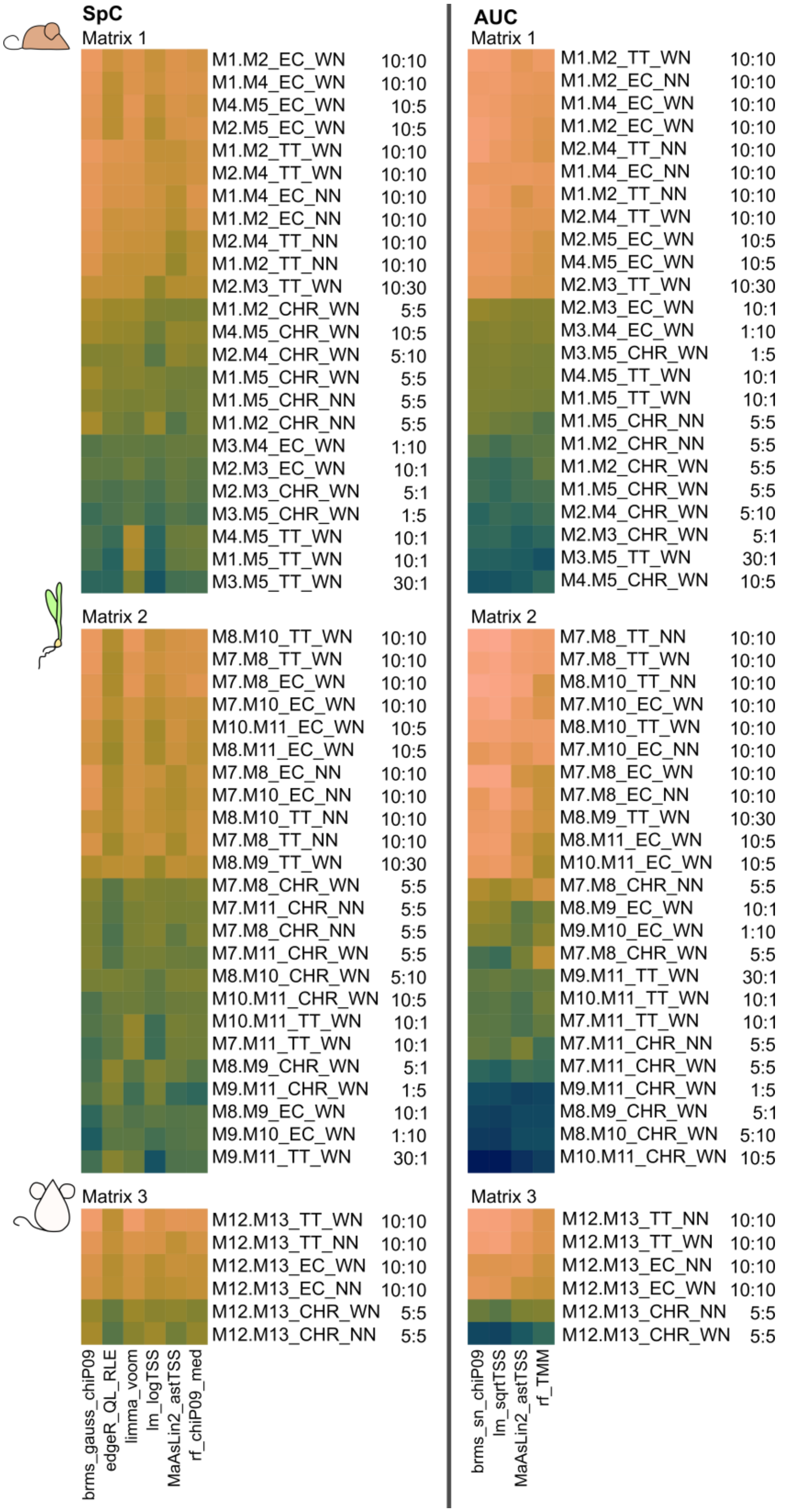


*Supplementary Figure 2: P4 scores for the best-performing tests of SpC and AUC comparisons per comparison. WN: with species-level normalization, NN: without species-level normalization.*

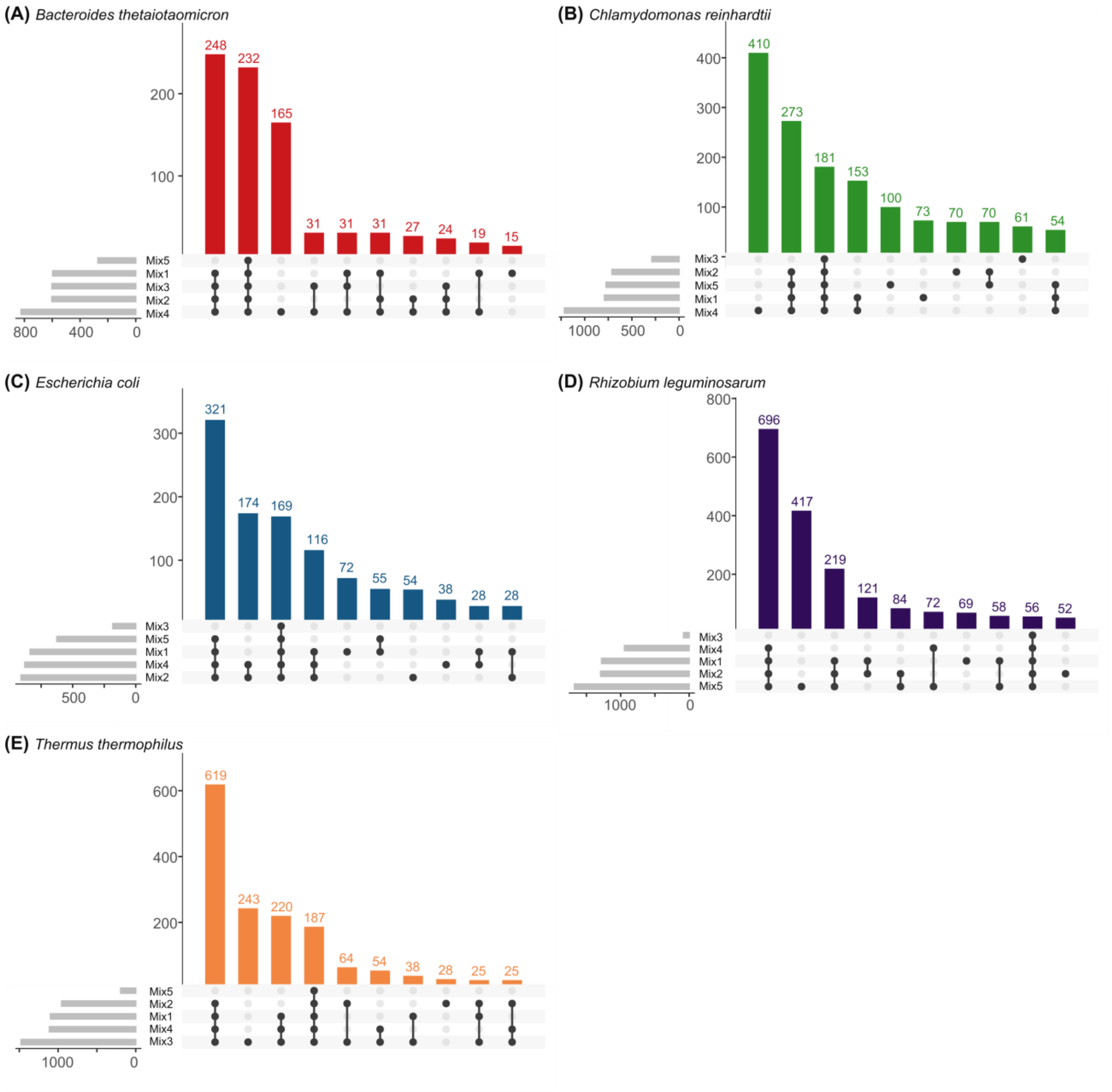


*Supplementary Figure 3: Numbers of protein groups identified per pure culture organism and mix, or combination of mixes, based on spectral count (SpC) data in the mouse faecal pellet matrix.*

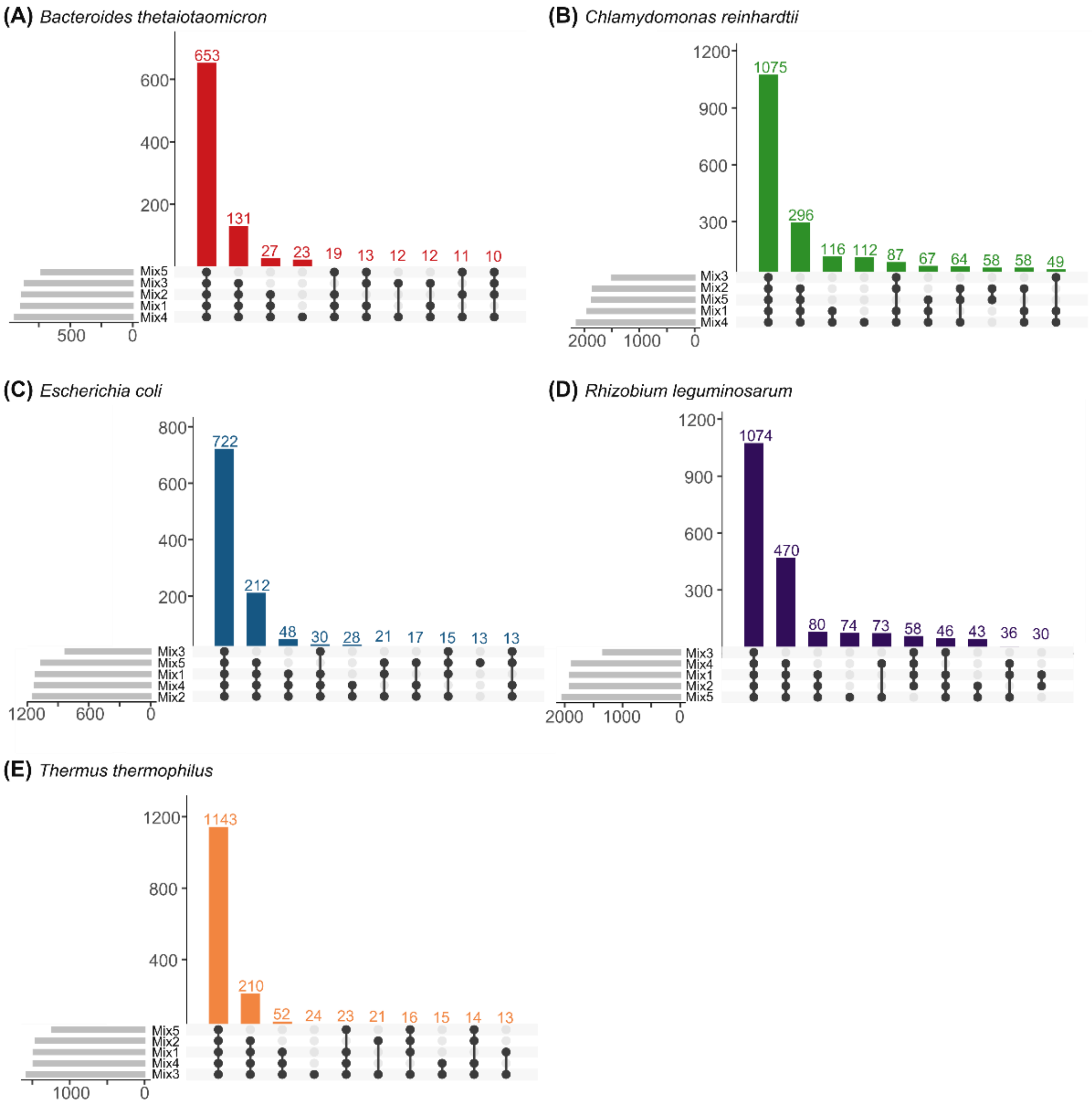


*Supplementary Figure 4: Numbers of protein groups identified per pure culture organism and mix, or combination of mixes, based on peptide intensity (AUC) data in the mouse faecal pellet matrix.*

#### Confounding effects decrease significant results

Comparisons of species (EC, TT, CRH) from M1 to M5 with the same species in M6 (Figure 1), where these species were mixed in from two culturing conditions (i.e., where these two conditions are confounded), generally decreased test performance, with the exception of NPV and TNR, which can increase if fewer proteins are detected as significantly differentially abundant overall. Additionally, AUC-based tests fared better overall on TPR as compared to SpC-based tests (Supplementary Figure 5). This result highlights that in many real-life datasets, e.g., biofilms with several phenotypes of the same organism being present [81], decreases in performance of the statistics have to be expected – meaning that only relatively stronger changes of expressed phenotypes can be accessed, if not accounting explicitly for spatial variation in the experimental design. There is no straightforward statistical approach to disentangle small changes in protein abundances and the presence of subpopulations with differing changes in protein abundances. Theoretically, changes in the protein abundances in different subpopulations can cancel each other out. Therefore, enrichment of sub-populations (e.g.,[82, 83]) or activity-based labeling such as stable isotope probing [84, 85] should be considered in order to draw meaningful conclusions about samples where sub-populations are to be expected.

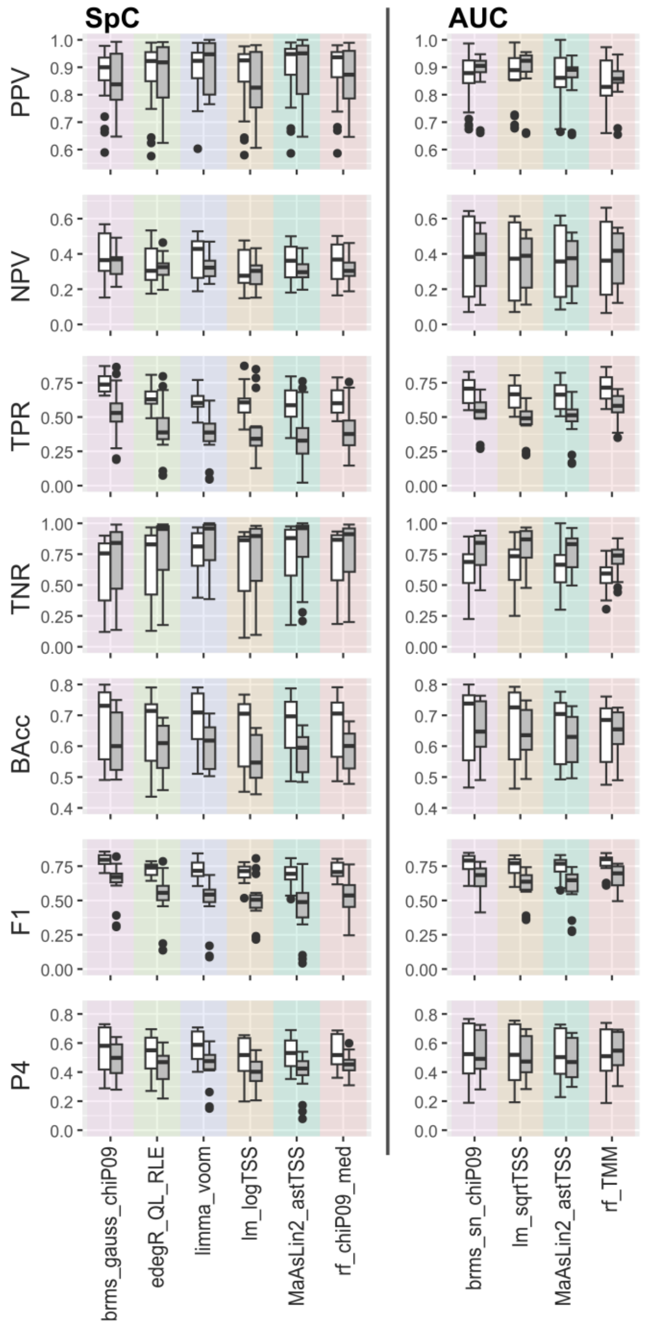


*Supplementary Figure 5: Comparison of test statistics for samples in which each species is only present from one culturing condition (white bars, M1 to M5 compared against each other for three species TT, EC, CRH) vs. a species present in a mix of two culture conditions (grey bars, M1 to M5 compared with EC, TT, CRH in M6). SpC: Spectral count quantification data; AUC: Area under the curve quantification data.*

#### Changing the ground truth test does not impact the overall pattern

We determined the extent to which our results are impacted by the choice of ground truth test by comparing the overlap between significant proteins identified in the pure cultures for several well-performing tests (Supplementary Figure 6). According to these, for SpC, our chosen test (brms_gauss_chiP09) identified slightly more proteins as significant (7-13% uniquely identified as significant) as compared to the other tests, indicating that overall there is agreement between tests on a large number of the significant proteins. For AUC, our chosen test lm_vsn was in accordance with brms_gauss_vsn and limma_vsn, whereas rf_TMM identified more protein groups as significant.

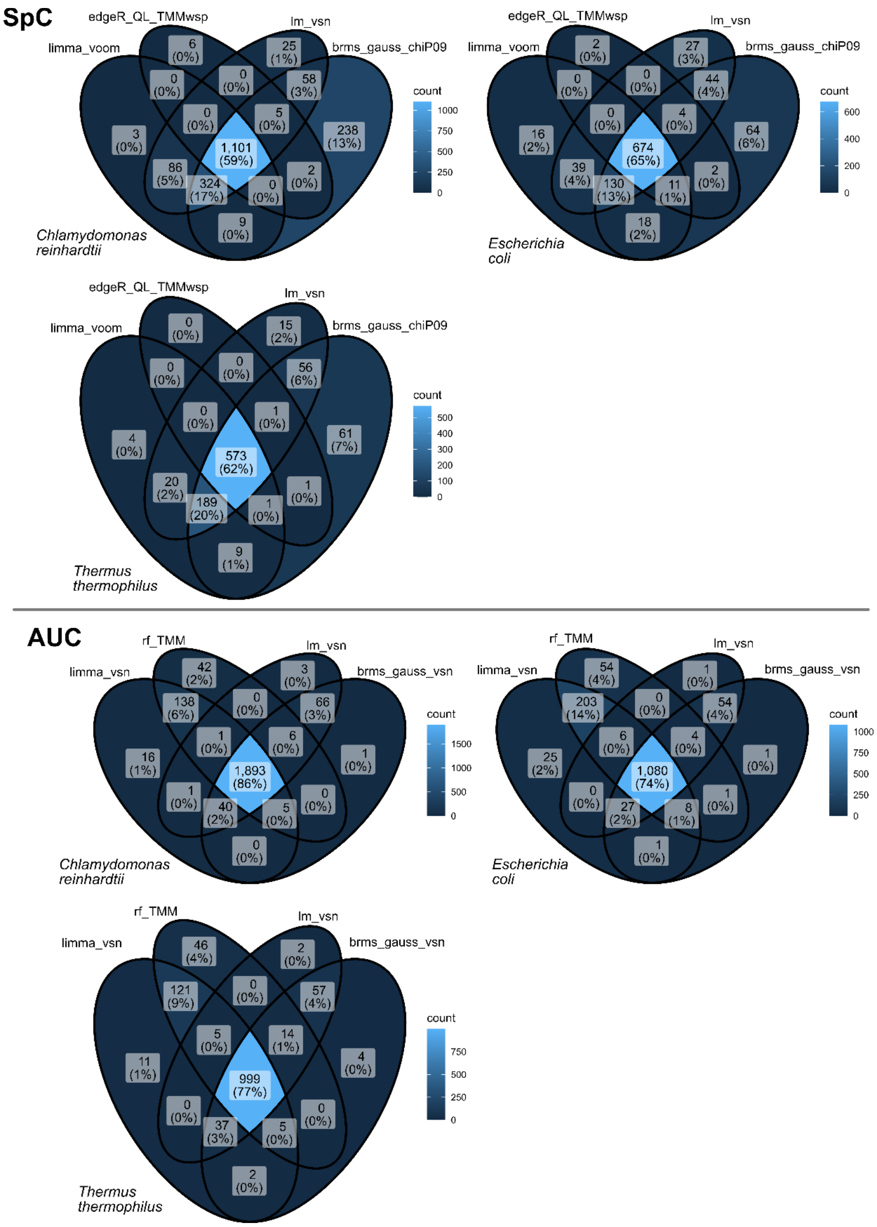


*Supplementary Figure 6: Overlap of different ground-truth tests in the identification of significant protein groups in the specified pure cultures. SpC: Spectral count quantification data; AUC: Area under the curve quantification data.*

Subsequently, we evaluated the overall pattern of statistical test performance based on a changed ground truth. For that, we decided on the test giving the most dissimilar results as compared to our ground truth test: For SpC, we used edgeR_QL_TMMwsp, which is more conservative as compared to brms_gauss_chiP09; for AUC, we used rf_TMM, which is more lenient as compared to lm_vsn. Based on the changed ground truths, the overall pattern of well-performing tests (i.e., tests with a median and lower quartile TNR and TPR in the top 25% of all tests) and some poorly-performing ones stayed the same (Supplementary Figure 7), highlighting the robustness of our approach. For SpC, the original ground-truth test (brms_gauss_chiP09) now moved out of the well-performing tests, due to edgeR_QL_TMMwsp being more conservative as compared to brms_gaus_chiP09, thus leading to a decreased TNR of brms_gauss_chiP09, and due to the overlap of proteins identified as significant in both methods being comparatively limited with about 60%. For AUC on the other hand, the original ground truth lm_vsn stayed within the list of well-performing tests, mainly because overlap between proteins identified as significant in the two ground truths was higher than for the compared SpC tests at 74-86%. We want to stress that the changed ground truth represents a compromise in performance (i.e., lower %TN/%TP) as compared to the ground truth used in the main analyses (see Main Results and Figure 3).

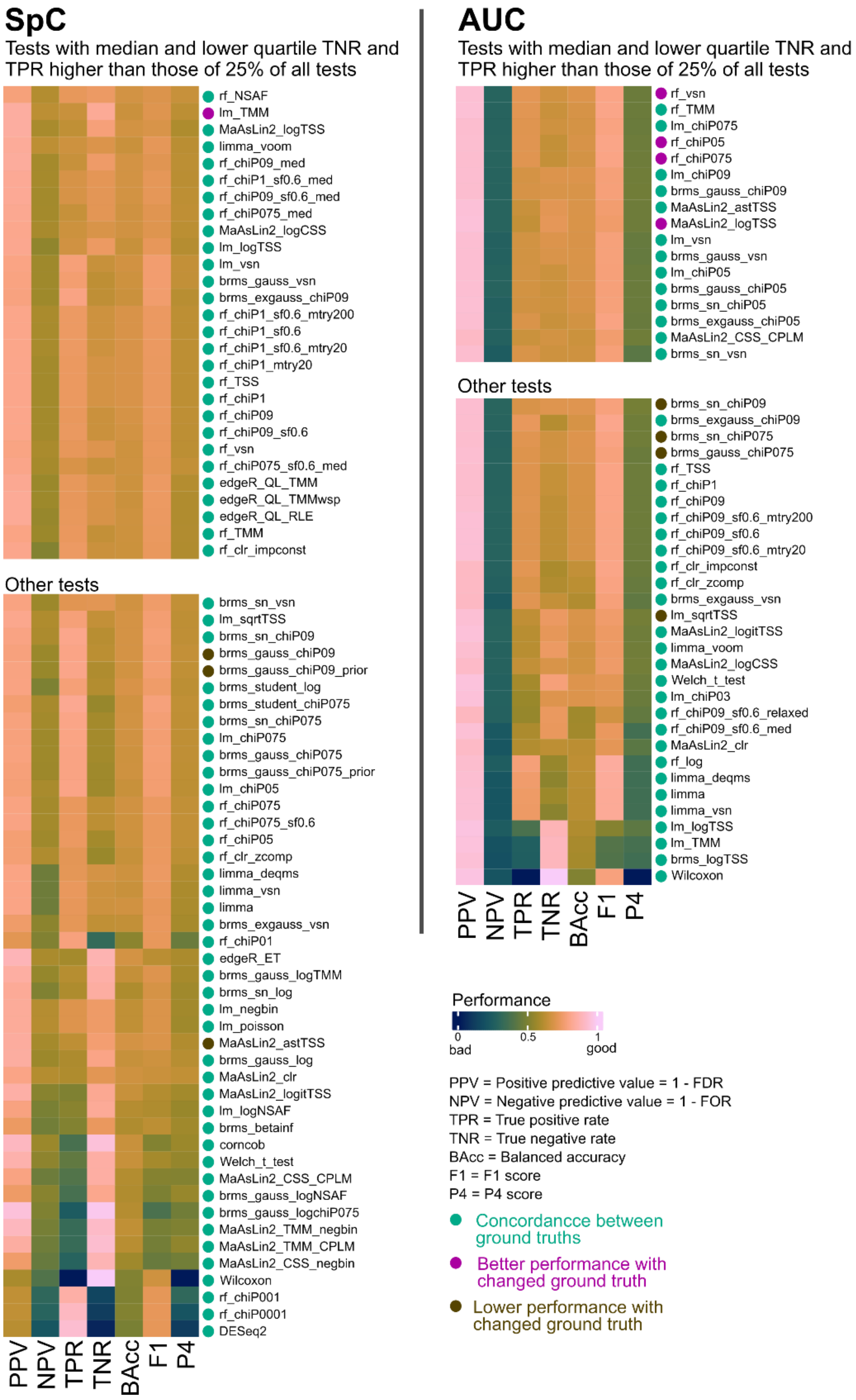


*Supplementary Figure 7: Median test evaluation metrics for mouse faecal pellet matrix comparisons (M1 to M5) for a changed ground truth test (SpC: edgeR_QL_TMMwsp; AUC: rf_TMM). Upper part: tests with a median and lower quartile TNR and TPR higher than those of 25% of all tests. Data was clustered row-wise based on Canberra distances. Colored dots next to the test names indicate whether the test was part of the well-performing tests or less well-performing tests as compared to the analyses based on the ground truth used in the main manuscript (see Main Text), or whether shifts occurred based on change of ground truth test. SpC: Spectral count quantification data; AUC: Area under the curve quantification data.*

### References

1. Bertani G. Studies on Lysogenesis. I. The mode of phage liberation by lysogenic Escherichia coli. J Bacteriol. 1951;62:293–300. https://doi.org/10.1128/jb.62.3.293-300.1951.

2. Harwood CR, Cutting SM. Chemically defined growth media and supplements. In: Molecular Biological Methods for Bacillus. 1990. p. 548.

3. Bacic MK, Smith CJ. Laboratory Maintenance and Cultivation of *Bacteroides* Species. Curr Protoc Microbiol. 2008;9. https://doi.org/10.1002/9780471729259.mc13c01s9.

4. Gorman DS, Levine RP. Cytochrome f and plastocyanin: their sequence in the photosynthetic electron transport chain of Chlamydomonas reinhardi. Proc Natl Acad Sci. 1965;54:1665–9. https://doi.org/10.1073/pnas.54.6.1665.

5. Vincent JM. A manual for the practical study of the root-nodule bacteria. Blackwell Scientific Publications, Oxford.; 1970.

6. Li W, Godzik A. Cd-hit: a fast program for clustering and comparing large sets of protein or nucleotide sequences. Bioinformatics. 2006;22:1658–9. https://doi.org/10.1093/bioinformatics/btl158.

7. Blakeley-Ruiz JA, Bartlett A, McMillan AS, Awan A, Walsh MV, Meyerhoffer AK, et al. Dietary protein source alters gut microbiota composition and function. ISME J. 2025;19:wraf048. https://doi.org/10.1093/ismejo/wraf048.

8. Wiśniewski JR, Zougman A, Nagaraj N, Mann M. Universal sample preparation method for proteome analysis. Nat Methods. 2009;6:359–62. https://doi.org/10.1038/nmeth.1322.

9. Parnell JJ, Pal G, Awan A, Vintila S, Houdinet G, Hawkes CV, et al. Effective Seed Sterilization Methods Require Optimization Across Maize Genotypes. Phytobiomes J. 2024;8:418–24. https://doi.org/10.1094/PBIOMES-12-23-0137-R.

10. Mordant A, Blakeley-Ruiz JA, Kleiner M. Metaproteomics-based stable isotope fingerprinting links intestinal bacteria to their carbon source and captures diet-induced substrate switching. ISME J. 2025;19:wraf127. https://doi.org/10.1093/ismejo/wraf127.

11. Bolger AM, Lohse M, Usadel B. Trimmomatic: a flexible trimmer for Illumina sequence data. Bioinformatics. 2014;30:2114–20. https://doi.org/10.1093/bioinformatics/btu170.

12. Li D, Liu C-M, Luo R, Sadakane K, Lam T-W. MEGAHIT: an ultra-fast single-node solution for large and complex metagenomics assembly via succinct *de Bruijn* graph. Bioinformatics. 2015;31:1674–6. https://doi.org/10.1093/bioinformatics/btv033.

13. Li H, Durbin R. Fast and accurate short read alignment with Burrows–Wheeler transform. Bioinformatics. 2009;25:1754–60. https://doi.org/10.1093/bioinformatics/btp324.

14. Kang DD, Li F, Kirton E, Thomas A, Egan R, An H, et al. MetaBAT 2: an adaptive binning algorithm for robust and efficient genome reconstruction from metagenome assemblies. PeerJ. 2019;7:e7359. https://doi.org/10.7717/peerj.7359.

15. Wu Y-W, Simmons BA, Singer SW. MaxBin 2.0: an automated binning algorithm to recover genomes from multiple metagenomic datasets. Bioinformatics. 2016;32:605–7. https://doi.org/10.1093/bioinformatics/btv638.

16. Pan S, Zhu C, Zhao X-M, Coelho LP. A deep siamese neural network improves metagenome-assembled genomes in microbiome datasets across different environments. Nat Commun. 2022;13:2326. https://doi.org/10.1038/s41467-022-29843-y.

17. Hickl O, Queirós P, Wilmes P, May P, Heintz-Buschart A. *binny* : an automated binning algorithm to recover high-quality genomes from complex metagenomic datasets. Brief Bioinform. 2022;23:bbac431. https://doi.org/10.1093/bib/bbac431.

18. Liu C-C, Dong S-S, Chen J-B, Wang C, Ning P, Guo Y, et al. MetaDecoder: a novel method for clustering metagenomic contigs. Microbiome. 2022;10:46. https://doi.org/10.1186/s40168-022-01237-8.

19. Sieber CMK, Probst AJ, Sharrar A, Thomas BC, Hess M, Tringe SG, et al. Recovery of genomes from metagenomes via a dereplication, aggregation and scoring strategy. Nat Microbiol. 2018;3:836–43. https://doi.org/10.1038/s41564-018-0171-1.

20. Hyatt D, Chen G-L, LoCascio PF, Land ML, Larimer FW, Hauser LJ. Prodigal: prokaryotic gene recognition and translation initiation site identification. BMC Bioinformatics. 2010;11:119. https://doi.org/10.1186/1471-2105-11-119.

21. Nesvizhskii AI, Aebersold R. Interpretation of Shotgun Proteomic Data. Mol Cell Proteomics. 2005;4:1419–40. https://doi.org/10.1074/mcp.R500012-MCP200.

22. Van Den Bossche T, Armengaud J, Benndorf D, Blakeley-Ruiz JA, Brauer M, Cheng K, et al. The microbiologist's guide to metaproteomics. iMeta. 2025;4:e70031. https://doi.org/10.1002/imt2.70031.

23. Palarea-Albaladejo J, Martín-Fernández JA. zCompositions — R package for multivariate imputation of left-censored data under a compositional approach. Chemom Intell Lab Syst. 2015;143:85–96. https://doi.org/10.1016/j.chemolab.2015.02.019.

24. Kleiner M. Normalization of metatranscriptomic and metaproteomic data for differential gene expression analyses: The importance of accounting for organism abundance. 2017. https://doi.org/10.7287/peerj.preprints.2846v1.

25. Mueller RS, Denef VJ, Kalnejais LH, Suttle KB, Thomas BC, Wilmes P, et al. Ecological distribution and population physiology defined by proteomics in a natural microbial community. Mol Syst Biol. 2010;6:374. https://doi.org/10.1038/msb.2010.30.

26. Zybailov B, Mosley AL, Sardiu ME, Coleman MK, Florens L, Washburn MP. Statistical Analysis of Membrane Proteome Expression Changes in *Saccharomyces cerevisiae*. J Proteome Res. 2006;5:2339–47. https://doi.org/10.1021/pr060161n.

27. Robinson MD, Oshlack A. A scaling normalization method for differential expression analysis of RNA-seq data. Genome Biol. 2010;11:R25. https://doi.org/10.1186/gb-2010-11-3-r25.

28. Chen Y, Chen L, Lun ATL, Baldoni PL, Smyth GK. edgeR v4: powerful differential analysis of sequencing data with expanded functionality and improved support for small counts and larger datasets. Nucleic Acids Res. 2025;53:gkaf018. https://doi.org/10.1093/nar/gkaf018.

29. Anders S, Huber W. Differential expression analysis for sequence count data. Genome Biol. 2010;11:R106. https://doi.org/10.1186/gb-2010-11-10-r106.

30. Law CW, Chen Y, Shi W, Smyth GK. voom: precision weights unlock linear model analysis tools for RNA-seq read counts. Genome Biol. 2014;15:R29. https://doi.org/10.1186/gb-2014-15-2-r29.

31. Zhu Y, Orre LM, Zhou Tran Y, Mermelekas G, Johansson HJ, Malyutina A, et al. DEqMS: A Method for Accurate Variance Estimation in Differential Protein Expression Analysis. Mol Cell Proteomics. 2020;19:1047–57. https://doi.org/10.1074/mcp.TIR119.001646.

32. Huber W, Von Heydebreck A, Sültmann H, Poustka A, Vingron M. Variance stabilization applied to microarray data calibration and to the quantification of differential expression. Bioinformatics. 2002;18 suppl_1:S96–104. https://doi.org/10.1093/bioinformatics/18.suppl_1.S96.

33. Legendre P, Gallagher ED. Ecologically meaningful transformations for ordination of species data. Oecologia. 2001;129:271–80. https://doi.org/10.1007/s004420100716.

34. Aitchison J. The Statistical Analysis of Compositional Data. J R Stat Soc Ser B Stat Methodol. 1982;44:139–60. https://doi.org/10.1111/j.2517-6161.1982.tb01195.x.

35. Greenacre M. The chiPower transformation: a valid alternative to logratio transformations in compositional data analysis. Adv Data Anal Classif. 2024;18:769–96. https://doi.org/10.1007/s11634-024-00600-x.

36. Bates D, Mächler M, Bolker B, Walker S. Fitting Linear Mixed-Effects Models Using **lme4**. J Stat Softw. 2015;67. https://doi.org/10.18637/jss.v067.i01.

37. Bürkner P-C. Advanced Bayesian Multilevel Modeling with the R Package brms. R J. 2018;10:395. https://doi.org/10.32614/RJ-2018-017.

38. Martin BD, Witten D, Willis AD. Modeling microbial abundances and dysbiosis with beta-binomial regression. Ann Appl Stat. 2020;14. https://doi.org/10.1214/19-AOAS1283.

39. Martin B, Witten D, Willis A. corncob: Count Regression for Correlated Observations with the Beta-Binomial. R package version 0.3.1. 2022.

40. Love MI, Huber W, Anders S. Moderated estimation of fold change and dispersion for RNA-seq data with DESeq2. Genome Biol. 2014;15:550. https://doi.org/10.1186/s13059-014-0550-8.

41. Ritchie ME, Phipson B, Wu D, Hu Y, Law CW, Shi W, et al. limma powers differential expression analyses for RNA-sequencing and microarray studies. Nucleic Acids Res. 2015;43:e47–e47. https://doi.org/10.1093/nar/gkv007.

42. Mallick H, Rahnavard A, McIver LJ, Ma S, Zhang Y, Nguyen LH, et al. Multivariable association discovery in population-scale meta-omics studies. PLOS Comput Biol. 2021;17:e1009442. https://doi.org/10.1371/journal.pcbi.1009442.

43. Breiman L. Random Forests. Mach Learn. 2001;45:5–32.

44. Janitza S, Celik E, Boulesteix A-L. A computationally fast variable importance test for random forests for high-dimensional data. Adv Data Anal Classif. 2018;12:885–915. https://doi.org/10.1007/s11634-016-0276-4.

45. Welch BL. The Generalization of `Student's' Problem when Several Different Population Variances are Involved. Biometrika. 1947;34:28. https://doi.org/10.2307/2332510.

46. Delacre M, Lakens D, Leys C. Why Psychologists Should by Default Use Welch's t-test Instead of Student's t-test. Int Rev Soc Psychol. 2017;30:92–101. https://doi.org/10.5334/irsp.82.

47. Benjamini Y, Hochberg Y. On the Adaptive Control of the False Discovery Rate in Multiple Testing with Independent Statistics. J Educ Behav Stat. 2000;25:60–83.

48. Benjamini Y, Yekutieli D. The control of the false discovery rate in multiple testing under dependency. Ann Stat. 2001;29:1165–88.

49. Kassambara A. rstatix: Pipe-Friendly Framework for Basic Statistical Tests. R package version 0.7.2. 2023.

50. Wilcoxon F. Individual Comparisons by Ranking Methods. Biom Bull. 1945;1:80. https://doi.org/10.2307/3001968.

51. Chen T, Guestrin C. XGBoost: A Scalable Tree Boosting System. In: Proceedings of the 22nd ACM SIGKDD International Conference on Knowledge Discovery and Data Mining. San Francisco California USA: ACM; 2016. p. 785–94. https://doi.org/10.1145/2939672.2939785.

52. Fernandes AD, Reid JN, Macklaim JM, McMurrough TA, Edgell DR, Gloor GB. Unifying the analysis of high-throughput sequencing datasets: characterizing RNA-seq, 16S rRNA gene sequencing and selective growth experiments by compositional data analysis. Microbiome. 2014;2:15. https://doi.org/10.1186/2049-2618-2-15.

53. Calgaro M, Romualdi C, Waldron L, Risso D, Vitulo N. Assessment of statistical methods from single cell, bulk RNA-seq, and metagenomics applied to microbiome data. Genome Biol. 2020;21:191. https://doi.org/10.1186/s13059-020-02104-1.

54. Nixon MP, Gloor GB, Silverman JD. Incorporating scale uncertainty in microbiome and gene expression analysis as an extension of normalization. Genome Biol. 2025;26:139. https://doi.org/10.1186/s13059-025-03609-3.

55. Gloor GB, Nixon MP, Silverman JD. Explicit Scale Simulation for analysis of RNA-sequencing count data with ALDEx2. NAR Genomics Bioinforma. 2025;7:lqaf108. https://doi.org/10.1093/nargab/lqaf108.

56. R Core Team. R: A Language and Environment for Statistical Computing. R Found Stat Comput Vienna Austria. 2023.

57. Wickham H, François R, Henry L, Müller K, Vaughan D. dplyr: A Grammar of Data Manipulation. R package version 1.1.4. 2023.

58. Wickham H. The Split-Apply-Combine Strategy for Data Analysis. J Stat Softw. 2011;40. https://doi.org/10.18637/jss.v040.i01.

59. Wickham H, Vaughan D, Girlich M. tidyr: Tidy Messy Data. R package version 1.3.1. 2024.

60. Wickham H, Averick M, Bryan J, Chang W, McGowan L, François R, et al. Welcome to the Tidyverse. J Open Source Softw. 2019;4:1686. https://doi.org/10.21105/joss.01686.

61. Gabry J, Češnovar R, Johnson A, Bronder S. cmdstanr: R Interface to "CmdStan". R package version 0.8.1. 2024.

62. van den Boogaart K, Tolosana-Delgado R, Bren M. compositions: Compositional Data Analysis. R package version 2.0-6. 2023.

63. Gamer M, Lemon J, Fellows I, Singh P. irr: Various Coefficients of Interrater Reliability and Agreement. R package version 0.84.1. 2019.

64. Kuznetsova A, Brockhoff PB, Christensen RHB. **lmerTest** Package: Tests in Linear Mixed Effects Models. J Stat Softw. 2017;82. https://doi.org/10.18637/jss.v082.i13.

65. Koncevičius K. matrixTests: Fast Statistical Hypothesis Tests on Rows and Columns of Matrices. R package version 0.2.3. 2023.

66. Wright MN, Ziegler A. ranger: A Fast Implementation of Random Forests for High Dimensional Data in *C++* and *R*. J Stat Softw. 2017;77. https://doi.org/10.18637/jss.v077.i01.

67. Magnusson A, Millar C. TAF: Transparent Assessment Framework for Reproducible Research. R package version 4.2.0. 2023.

68. Kuhn M, Wickham H. Tidymodels: a collection of packages for modeling and machine learning using tidyverse principles. 2020. https://www.tidymodels.org.

69. Microsoft Corporation, Weston S. doParallel: Foreach Parallel Adaptor for the "parallel" Package. R package version 1.0.17. 2022.

70. Microsoft, Weston S. foreach: Provides Foreach Looping Construct. R package version 1.5.2. 2022.

71. Gu Z, Gu L, Eils R, Schlesner M, Brors B. *circlize* implements and enhances circular visualization in R. Bioinformatics. 2014;30:2811–2. https://doi.org/10.1093/bioinformatics/btu393.

72. Gu Z. Complex heatmap visualization. iMeta. 2022;1:e43. https://doi.org/10.1002/imt2.43.

73. Auguie B. egg: Extensions for "ggplot2": Custom Geom, Custom Themes, Plot Alignment, Labelled Panels, Symmetric Scales, and Fixed Panel Size. R package version 0.4.5. 2019.

74. Tiedemann F. gghalves: Compose Half-Half Plots Using Your Favourite Geoms. R package version 0.1.4. 2022.

75. Wickham H. ggplot2: Elegant Graphics for Data Analysis. 2nd ed. 2016. Cham: Springer International Publishing : Imprint: Springer; 2016.

76. Pedersen T. patchwork: The Composer of Plots. R package version 1.1.3. 2023.

77. Neuwirth E. RColorBrewer: ColorBrewer Palettes. R package version 1.1-3. 2022.

78. Pedersen T, Crameri F. scico: Colour Palettes Based on the Scientific Colour-Maps. R package version 1.5.0. 2023.

79. Crameri F, Shephard GE, Heron PJ. The misuse of colour in science communication. Nat Commun. 2020;11:5444. https://doi.org/10.1038/s41467-020-19160-7.

80. Wickham H, Henry L, Pedersen T, Luciani T, Decorde M, Lise V. svglite: An “SVG” Graphics Device. R package version 2.1.3. 2023.

81. Stewart PS, Franklin MJ. Physiological heterogeneity in biofilms. Nat Rev Microbiol. 2008;6:199–210. https://doi.org/10.1038/nrmicro1838.

82. Babin BM, Atangcho L, Van Eldijk MB, Sweredoski MJ, Moradian A, Hess S, et al. Selective Proteomic Analysis of Antibiotic-Tolerant Cellular Subpopulations in *Pseudomonas aeruginosa* Biofilms. mBio. 2017;8:e01593-17. https://doi.org/10.1128/mBio.01593-17.

83. Hinzke T, Kleiner M, Meister M, Schlüter R, Hentschker C, Pané-Farré J, et al. Bacterial symbiont subpopulations have different roles in a deep-sea symbiosis. eLife. 2021;10:e58371. https://doi.org/10.7554/eLife.58371.

84. Kleiner M, Kouris A, Violette M, D’Angelo G, Liu Y, Korenek A, et al. Ultra-sensitive isotope probing to quantify activity and substrate assimilation in microbiomes. Microbiome. 2023;11:24. https://doi.org/10.1186/s40168-022-01454-1.

85. Sachsenberg T, Herbst F-A, Taubert M, Kermer R, Jehmlich N, Von Bergen M, et al. MetaProSIP: Automated Inference of Stable Isotope Incorporation Rates in Proteins for Functional Metaproteomics. J Proteome Res. 2015;14:619–27. https://doi.org/10.1021/pr500245w.
